# Characterization of RNA Polymerase II Trigger Loop Mutations using Molecular Dynamics Simulations and Machine Learning

**DOI:** 10.1101/2022.08.11.503690

**Authors:** Bercem Dutagaci, Bingbing Duan, Chenxi Qiu, Craig D. Kaplan, Michael Feig

## Abstract

Catalysis and fidelity of multisubunit RNA polymerases rely on a highly conserved active site domain called the trigger loop (TL), which achieves roles in transcription through conformational changes and interaction with NTP substrates. The mutations of TL residues cause distinct effects on catalysis including hypo- and hyperactivity and altered fidelity. We applied molecular dynamics simulation (MD) and machine learning (ML) techniques to characterize TL mutations in the *Saccharomyces cerevisiae* RNA Polymerase II (Pol II) system. We did so to determine relationships between individual mutations and phenotypes and to associate phenotypes with MD simulated structural alterations. Using fitness values of mutants under various stress conditions, we modeled phenotypes along a spectrum of continual values. We found that ML could predict the phenotypes with 0.68 *R*^2^ correlation from amino acid sequences alone. It was more difficult to incorporate MD data to improve predictions from machine learning, presumably because MD data is too noisy and possibly incomplete to directly infer functional phenotypes. However, a variational auto-encoder model based on the MD data allowed the clustering of mutants with different phenotypes based on structural details. Overall, we found that lethal mutations tended to increase distances of TL residues to the NTP substrate, while viable loss-of-function (LOF) substitutions tended to confer an increase in distances between TL and bridge helix (BH). In contrast, GOF mutants generally have a disrupting effect on hydrophobic contacts among TL and nearby helices.

**AUTHOR SUMMARY:** RNA polymerase II (Pol II) synthesizes RNA with the help of an active site domain called trigger loop (TL). The mutations of TL cause changes in the activity of Pol II that could range from gain-of function (GOF) to loss-of-function (LOF) or lethal. This study provides a systematic characterization of the structural and functional outcomes of the TL mutations using molecular dynamics (MD) simulations and machine learning (ML). We obtained functional phenotypes of mutants by ML using the genetic fitness scores as the input. We revealed that mutant TL sequences could predict the functional outcomes at a relatively high correlation. Then, we performed MD simulations to relate the structural information to the phenotypes. The analysis of the MD data suggested that the lethal mutants had increased distances between the TL and the substrate, while a subset of LOF mutants showed increased distances between TL and another active site domain called bridge helix (BH). On the other hand, GOF mutants had effects on the hydrophobic interactions around the active site. Overall, this study enhances our understanding of the effects of TL mutations to the Pol II function.

## INTRODUCTION

RNA polymerase II (Pol II) is the enzyme that synthesizes mRNA in eukaryotes. Structural (1–4) and computational (5–8) studies have provided insights into the mechanism of this process, which takes place by repeating a nucleotide addition cycle (NAC) for addition of new nucleotides to the nascent RNA (9). Proposed mechanisms for the NAC emphasize conformational changes of a highly conserved domains in the active site, present within the largest subunit of yeast Pol II, Rpb1. One of these domains is called the trigger loop (TL) and the other is the nearby bridge helix (BH). The TL has open and closed conformations, which are known to be important for the nucleotide addition (10–13). The NAC starts with the Pol II complex with an open TL that allows an incoming nucleoside triphosphate (NTP) to enter the active site. Upon initial binding of the NTP, the TL closes and catalysis is promoted for substrates base-paired with the template. This results in the pre-translocation (substrate added) state, followed by TL opening together with the pyrophosphate ion (PPi) release, and subsequent or concurrent translocation (14). The TL has been suggested to have an important function in selecting (4, 15–17) and positioning (18, 19) the correct NTP at the active site and in affecting the kinetics (15, 20, 21) of the NAC. TL involvement during transcriptional pausing (22, 23), backtracking (24, 25) and translocation (10, 26, 27) has also been proposed.

Detailed mechanisms of TL function are still not fully known. In previous studies, a complete deletion of the TL from different species caused marked reductions in transcription rate (23, 28, 29). Certain residues were identified to be especially important for function, such as H1085, L1081, E1103 and Q1078 (4, 15, 16, 18, 26, 30–32). H1085 and L1081 are in close distance to the NTP when the TL is closed. Therefore, their roles have been attributed with the positioning of the correct NTP. Most of substitutions of H1085 and L1081 are lethal (4, 18, 32). On the other hand, E1103 mutations are known to cause an increased catalytic rate but with compromised fidelity (15, 16, 26, 32). Further studies showed that Q1078 has interactions with the sugar moiety of the NTP, and most of its mutations are also not viable (30–32). Because the TL must support multiple conformations, there may be complex effects of specific substitutions. Site-directed mutagenesis and a prior comprehensive genetic study on TL alleles together suggest Pol II mutant phenotypic classes and complex interactions between residues supporting a functional network (21, 32). In that study, the effects of mutations were classified broadly as either ‘loss of function’ (LOF), where catalytic activity is, or is predicted to be, reduced *in vitro*, ‘gain of function’ (GOF), where catalytic activity is, or is predicted to be, increased, or ‘lethal’, where essential functions are compromised. As a result, genetic phenotypes were associated different functional outcomes providing a framework for insights into the role of different TL residues during transcription.

How the functional TL phenotypes that result from residue variations are manifested is an open question, since structural and dynamic details at the atomic level for individual mutants is lacking, as is understanding of potential commonalities at the biochemical level within mutant classes. To address this, we combined the data from experimental fitness scores and molecular dynamics (MD) simulations for TLs with different amino acid sequences to predict functional and structural outcomes of TL mutations using machine learning (ML) frameworks. The analysis here is based on an updated fitness dataset that extends the earlier analysis of Qiu *et al*. to develop a complete TL mutation phenotype map based on a continuous representation of functional phenotypes (32). First, we developed ML models using amino acid sequences to predict TL mutation phenotypes. Then, we selected 135 TL single mutants with known functional phenotypes and performed atomistic molecular dynamics (MD) simulations of those mutants. Following MD simulations, we applied ML algorithms on data extracted from the simulations to develop a better understanding how different phenotypes map onto differences in structure and dynamics of Pol II near the active site. The structural data obtained from the MD simulations was primarily used to provide a mechanistic understanding of the TL mutant phenotypes when used in a variational auto-encoder (VAE) framework that allowed us to map function to structural features. Specific insights from this analysis are that lethal mutants have generally increased intramolecular distances between TL residues and the NTP, while a subset of the LOF mutants have large distances between TL and BH residues. We also predicted two distinct classes of GOF phenotypes where both affect a hydrophobic pocket formed by active site residues while a subset has increased BH-TL interactions. Overall, these findings lead to further understanding of the specific roles of the TL and the BH during Pol II function. This study also suggests that longer MD simulations, on possibly μs time scale, might be required to enhance the inference of the mutant mechanisms.

## RESULTS

This study focuses on the interpretation of Pol II TL mutation phenotypes based on experiments via MD simulations and ML to develop a deeper mechanistic understanding of the role of the TL during transcription. We describe here three sets of results: 1) Based on fitness data from second-generation deep mutational scanning of the Pol II TL in *S. cerevisiae*, we generated a model for the classification of TL mutants along a continuous phenotypic spectrum; 2) we applied ML to infer TL mutant phenotypes from TL sequence with and without structural data from MD simulations using a subset of gold standard mutants as training data; and, 3) we extracted mechanistic principles for how different classes of TL mutations modulate Pol II function based on variational auto-encoder (VAE) models that were trained on MD simulation data.

### Continuous Pol II function phenotypes for TL mutations from experimental fitness data

Previous deep mutational scanning of the Pol II TL in yeast had classified mutants as lethal, LOF, GOF, or indeterminate(32). In this study, we classified mutants based on a phenotypic numerical continuum instead of discrete classes. GOF, LOF, and lethal phenotypes were mapped to values of +1, −1 and −2, respectively and a value of 0 corresponds to indeterminate phenotypes or the WT. This allowed us to further project the phenotypes into a two-dimensional latent space and analyze the transition between phenotypes. To place individual mutants along a phenotypic continuum, we first obtained quantitative fitness data for all possible TL single substitution mutants and a range of selected double mutants for a number of selection conditions (Fig. 1A), (see Methods). These conditions have previously been shown to enable discrimination among hypo- and hyperactive Pol II mutants (LOF and GOF, respectively)(32). We trained a neural network model based on the fitness data against the annotated functional phenotypes, converted to their corresponding numerical values, for a subset of the mutations described in the previous study (3, 21, 32). The resulting ML model was then used to predict the functional phenotype from the fitness data for the complete set of mutations in the TL as described in the Methods (Fig. 1B). Most substitutions of residues between T1077 to G1088 were predicted as lethal or LOF consistent with the study of Qiu et al (32) and consistent with the fact that these TL residues are critical for function. Mutations of V1094, P1099, and to a lesser extent, L1105 broadly resulted in LOF phenotypes, suggesting importance for WT fitness. Conversely, substitutions of some other residues mostly result in GOF phenotypes, as previously suggested. Among those, A1076, M1079, G1097, and L1101 form a hydrophobic pocket that may stabilize the open TL state (30). Disruption of this pocket by mutations may facilitate TL closing leading to a GOF phenotype (30, 32), albeit a bias towards TL closing may come at the expense of decreased fidelity (15–17, 26, 33, 34). In addition, selected mutations at K1092, K1093, R1100 and most at E1103 lead to GOF phenotypes. The residues K1092 and K1093 were suggested to stabilize the TL open state through interactions with other Rpb1 residues like D716, and D1309/E1280, respectively, shown by simulation and experimental studies (12, 30, 32). Similarly, the GOF phenotype observed for E1103 substitutions was attributed to their stabilizing effect on the closed TL state (15, 21, 26).

**Figure 1.**
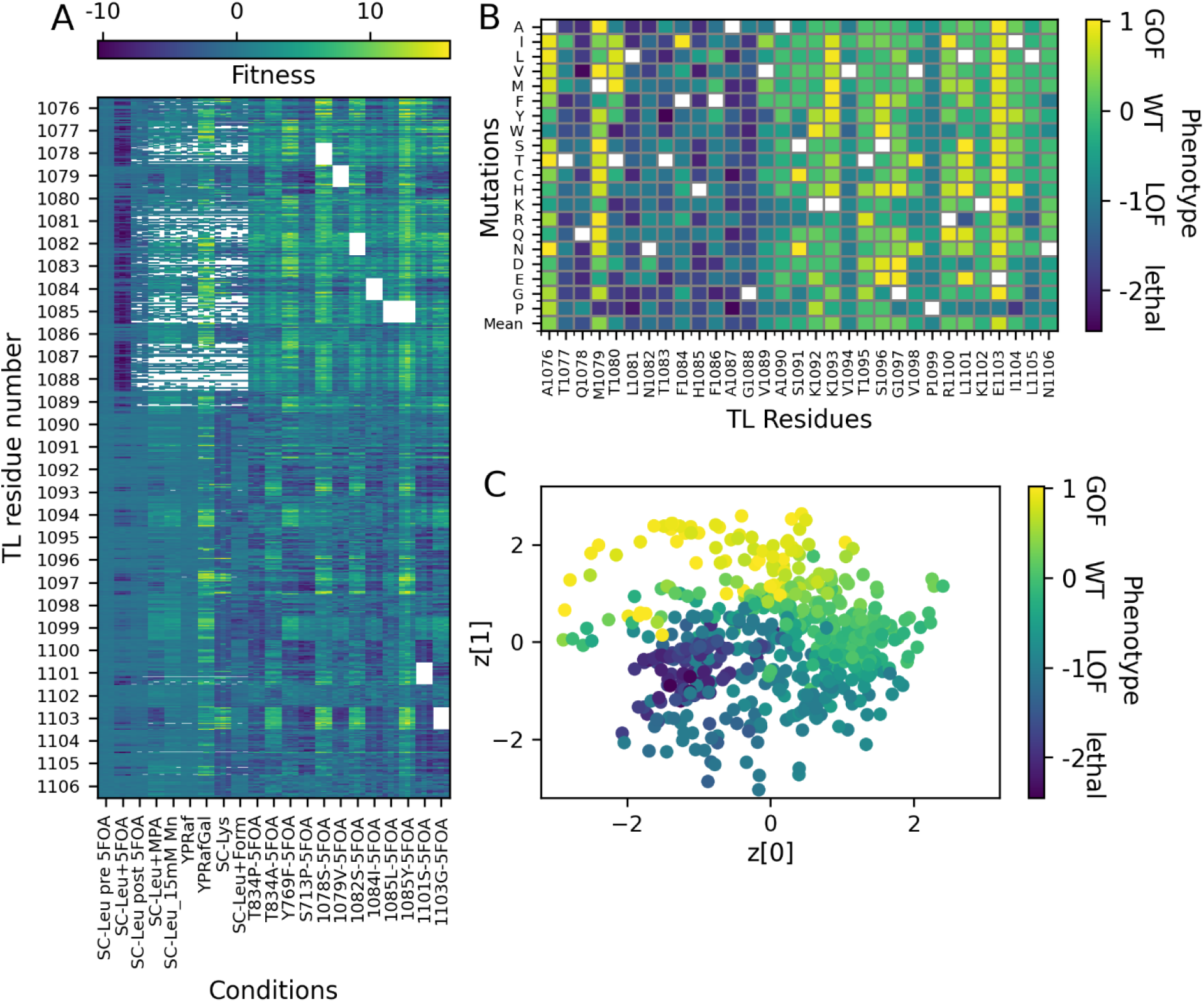
ML prediction of phenotypes from fitness data. (A) Fitness scores for mutants of TL residues under different conditions (see x-axis labels). For each residue, there are 21 data points with three replicates each; (B) phenotypical landscape predicted from the fitness data. Each box is colored with the phenotype; white boxes reflect the WT amino acid at those positions; the row at the bottom depicted as “Mean” shows the average phenotypes for the residues (C) latent space of the unsupervised VAE model based on the fitness data with each data point colored according to its corresponding phenotype.

To reduce dimensionality of the mutant space and associate the locations of the mutants with different phenotypes, we mapped the fitness values of the mutants onto a two-dimensional latent space by applying a VAE model. Fig. 1C shows the resulting latent space distribution of each mutant, colored according to the predicted phenotypes. Although continuous phenotype predictions benefitted from supervision based on known phenotypes, the VAE model was trained without such supervision. Nevertheless, there is clear clustering of the mutants according to the predicted phenotypes. The VAE model provides a regularized latent space with a more gradual transition of the phenotypes compared to the relatively more distinct locations of the different phenotypes in a 2D PCA analysis (Fig S1). Moreover, the gradual transition between different phenotype classes suggests that this type of classification provides additional insights that are not captured by a discrete classification and that may be more consistent with an evolutionary fitness landscape. Interestingly, transitions between nearby latent space projections are not just between neighboring phenotypes (*e*.*g*. from lethal to LOF and from LOF to neutral and then GOF) but also almost directly from lethal to GOF for some mutations (Fig. 1D). This suggests that phenotypes can be reversed with just a few mutations in some cases or alternatively, some substitutions turn to lethal because of manifesting a strong GOF phenotype.

### Inferring phenotypes from TL sequence

To further understand the information about fitness encoded in the TL sequence, we trained a supervised neural network to predict function from sequence. The target data were the continuous phenotype values determined from the experimental fitness data as described above (Supplementary Spreadsheet 1). In total, ten model replicates were generated for ten randomly generated training (100 mutants) and test (35 mutants) sets to result in 100 models in total. Then, we took the models that provided the best *R*^*2*^ and slope combination for each test set. The predictions from the best models for the ten sets were ensemble averaged to obtain the overall predictions. These are provided in the Supplementary Spreadsheet 1. The training and test loss of the ten models is shown in Fig. S2 and the correlations for the training and test sets for each model are shown in Figs. S3 and S4, respectively. Fig. 2A shows the average prediction performance with good correlation (*R*^2^ = 0.68). However, the slope of 0.60 and a more limited range in predicted values compared to the actual phenotypes indicates that extreme outcomes (gain of function or lethal) are not predicted as reliably as the overall trend. This is also evident in the difference map for predictions shown in Fig. S5. Fig. 2B shows sequence-based phenotype predictions of single mutations. The predictions largely agree with the phenotypes from the fitness values, but, again, with less variation in the predicted values towards the extremes. We also compared sequence-based model with a simple model that predicts the phenotypes from the average phenotypes for each mutant from the training sets used in the sequence models (see Fig S6). Compared to the sequence-based model we found a lower overall correlation (*R*^2^=0.59) but an improved slope (0.71). However, for the sequence model, there were difficulties in predicting phenotypes that have a large deviation from the average phenotype of each residue (see the outliers of M1079P, I1104P, L1101R, T1080M in Fig. S5). Overall, the improved correlation obtained by the sequence model over the simple model suggests that the TL sequence is a powerful predictive feature for inferring functional phenotypes of TL residue mutants.

**Figure 2.**
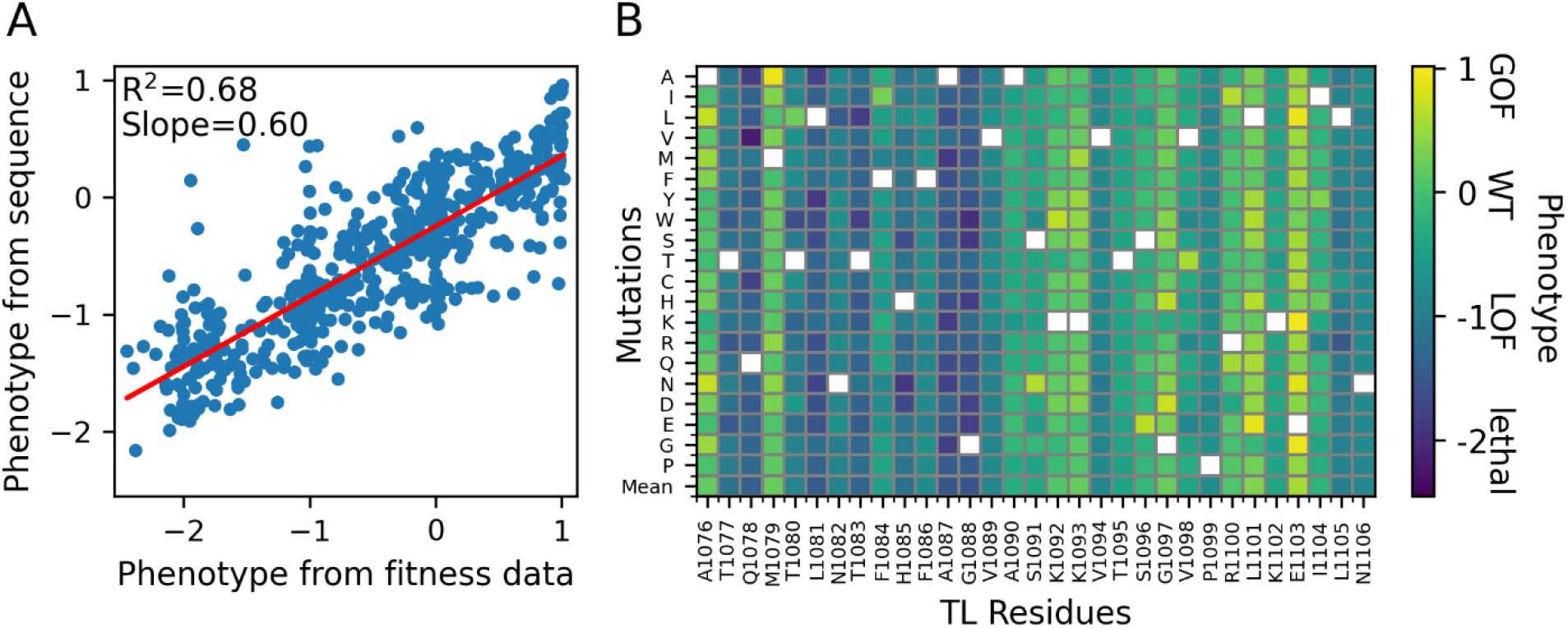
Prediction of phenotype from TL sequence. (A) Phenotype predicted from sequence vs. phenotype obtained from fitness data along with a linear regression curve. The predictions are the ensemble average values from models trained with 10 random training/test sets (B) Phenotypes predicted from the sequence for all single mutations.

We further tested the models trained on single mutants for the prediction of double mutants. We ensemble averaged the predictions of ten models as we did for the single mutants. Fig. 3 shows the phenotypic landscapes of all individual TL substitutions combined with either E1103G (GOF), G1097D (GOF), F1084I (GOF) or Q1078S (LOF). Generally, our model predicted additive effects of double mutants on phenotypes in which similar phenotypes showed an increase effect while opposite phenotypes suppressed each other. More specifically, the GOF mutants (E1103G, G1097D, F1084I) were predicted to cause the suppression of LOF and lethal mutants across the entire set of additional mutations. This additive prediction is consistent with previous studies(16, 21). Increased GOF phenotypes for GOF-GOF double mutants were also predicted. These predictions were in contrast to GOF combinations E1103G-G1097D and E1103G-F1084I having been found to be lethal(21). These combinations were hypothesized to be lethal due to extreme GOF phenotypes crossing a threshold for viability, an outcome not included in the training of ML model based on single mutations. The LOF mutant of Q1078S also predicted additive effects by suppressing the GOF phenotypes and causing lethal or more severe LOF phenotypes for LOF-LOF double mutants. We note that neutral (WT) function may be restored for certain double mutations of M1079, K1092, K1093, G1097, L1101, when added to Q1078S according to the predictions. It remains to be tested experimentally whether such double mutants could in fact restore normal function.

**Figure 3.**
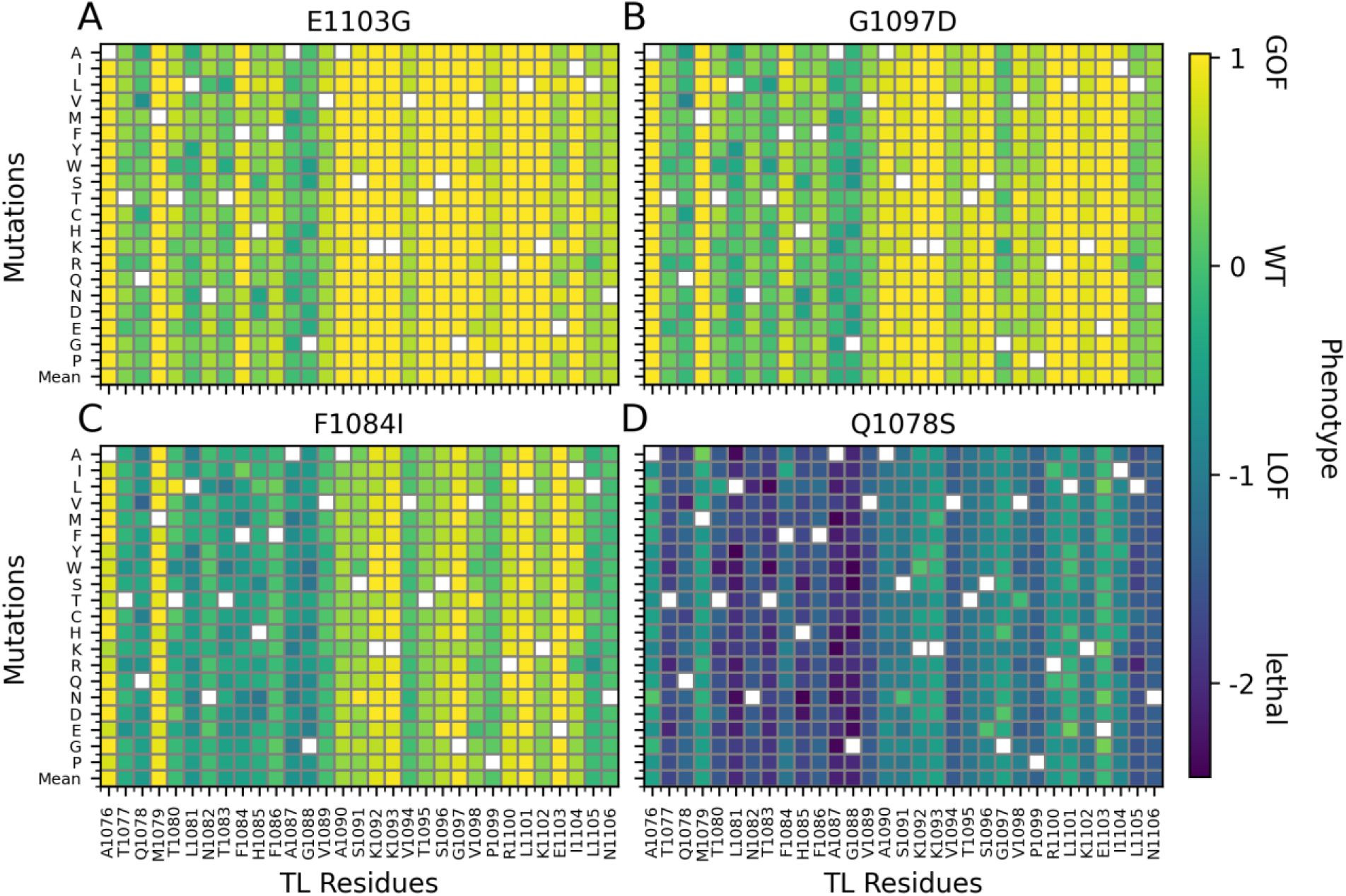
Prediction of phenotypes from TL sequence for double mutants of E1103G (A), G1097D (B), F1084I (C) and Q1078S (D).

### Inferring phenotypes from MD simulations of mutants

MD simulations were performed on 135 mutants to determine if emergent properties of the simulated mutants might provide better discrimination across mutant classes or between phenotypes, and, if so, what structural and dynamic properties might be hallmarks of specific phenotypical outcomes. Mutants were chosen based on the predictions from the previous studies (21, 32). To generate features for ML training, we extracted intramolecular distance data from the MD simulations (Fig. 4). Specifically, we captured a subset of intramolecular distances from the MD trajectories deemed to be relevant for Pol II function and potentially sensitive to TL mutations: TL-TL residue pairs, TL-BH residue pairs and BH-BH residue pairs and TL residue-GTP pairs that are at close distance in the WT structure; GTP P_α_ and terminal RNA O3’ distance relevant for catalysis; base pair distance between GTP-H1 and the corresponding DNA (18-DNA-N3) and the distance between sugar carbons of GTP-C1’ and 18-DNA-C1’; finally, the distance between Mg^2+^ and P_α_ of GTP. In total, 62 distances were calculated, and the distance lists and average distances are provided in the Supplementary Spreadsheet 2.

**Figure 4.**
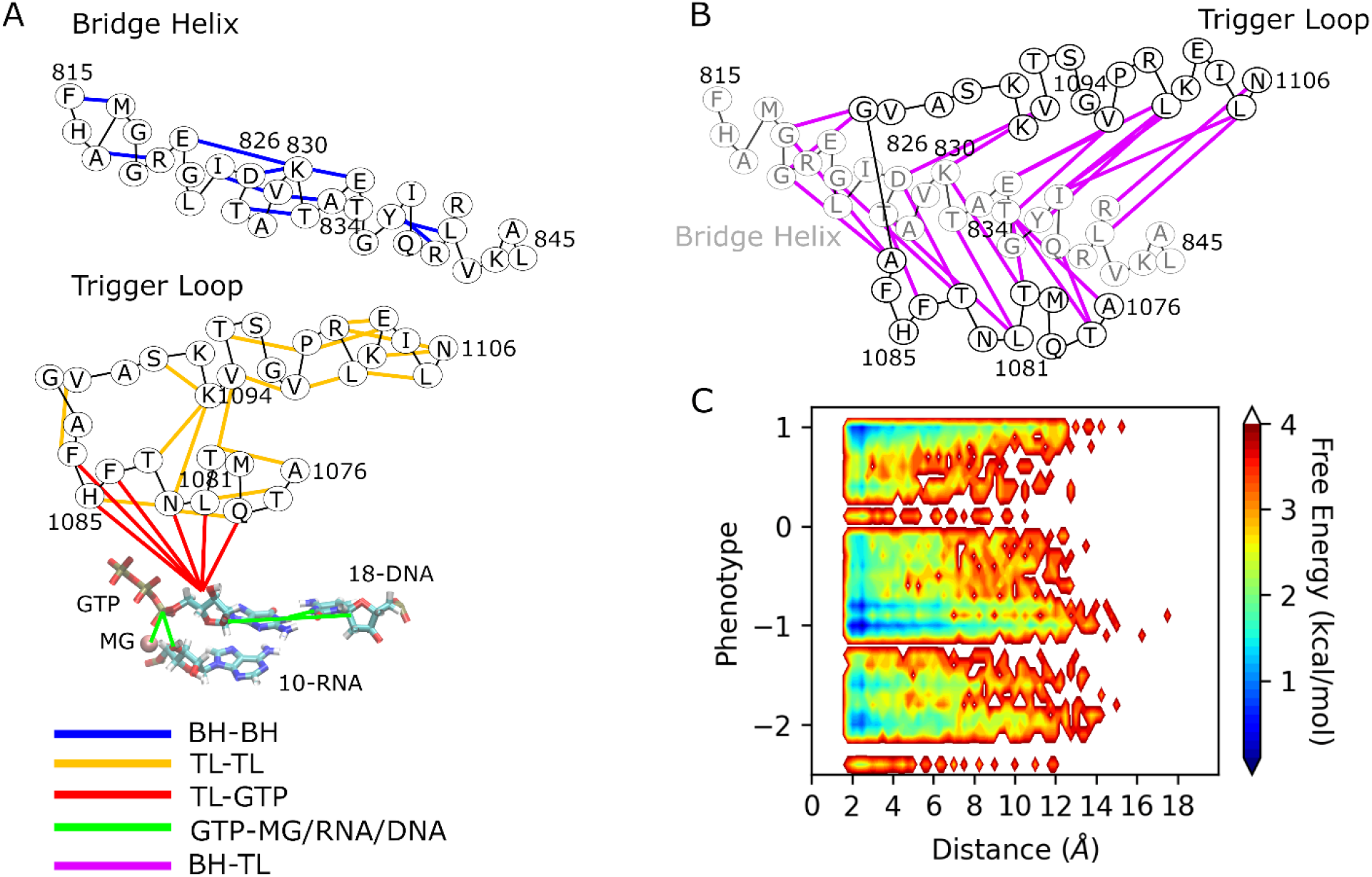
Distance analysis of the MD trajectories. (A) Schematic illustration of the pairs of BH-BH, TL-TL, TL-GTP and GTP with Mg^2+^, terminal RNA and base-pair DNA and (B) BH-TL with the color codes given in the figure, the black lines are used to show the adjacent amino acids (C) heatmap plot of phenotypes vs average distances for the mutants from MD simulations. Distances are provided in the Supplementary Spreadsheet 2.

Fig. 4-A and 4-B show the schematic representation of the distances and Fig 4-C shows the free energy map for the phenotype vs. average distances for each mutant studied via simulation. Most of the distances stay between 2-12 Å, while as the phenotypes range from GOF (1.0) to lethal (−2), some distances become longer. However, the affected distances differ between mutants. The lethal mutants (phenotype < −1.5) have effects mostly on the TL-GTP distances (Fig. S7). On the other hand, the distances between TL and BH residues are affected by mostly LOF mutants, those might be indirectly affecting the catalysis (Fig. S7). This overall suggests that lethal and LOF mutants cause increase in distances either of GTP or TL-BH residues that directly or indirectly impact the catalysis.

ML models for predicting phenotypes were then trained using the MD data with and without sequence data (Fig. 5, Figs. S2-4). ML models using only the MD-derived distances were clearly not as predictive as the models based on just amino acid sequence, even when a more complex model with an attention layer was considered (Fig. 5). We note that the *R*^2^ values in Fig. 2 and Fig. 5 are different since Fig 2 shows the correlations of the average predictions over the sets for all the mutants while Fig. 5 shows the averaged *R*^2^ over the sets for the mutants in the test sets. We also attempted to combine MD and sequence data and found that the MD data could not improve the predictions over using just the sequence information by itself. We interpret this finding to suggest that the MD data may be too noisy and sampling may be incomplete to reliably discern differences between specific mutations. On the other hand, knowledge about different amino acids implicitly contains information about amino acid sizes and physical characteristics such as charge and hydrophobicity. Taken together, this may be sufficient to characterize the effects of mutations with respect to phenotypic outcomes.

**Figure 5.**
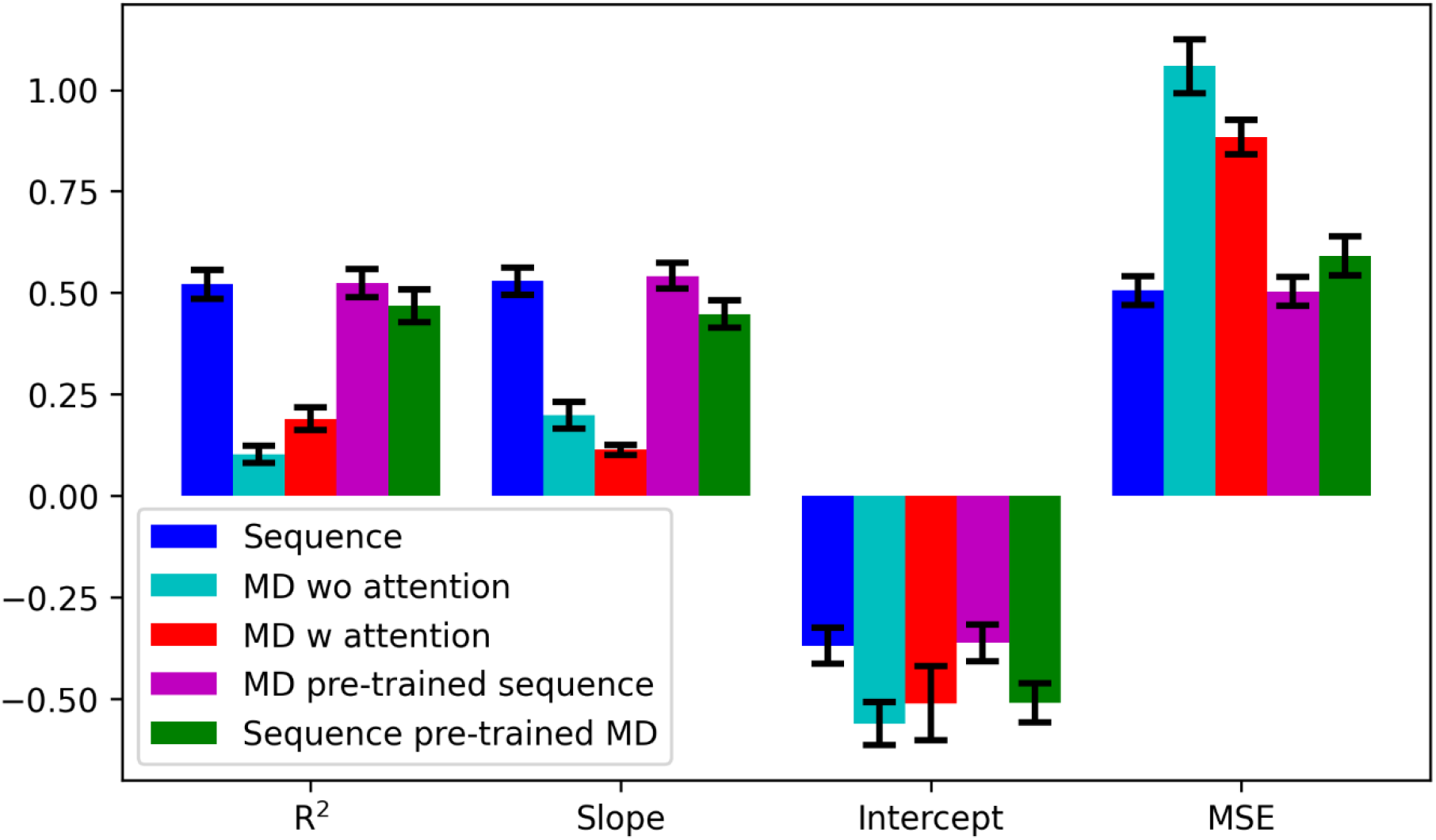
Performance of phenotype predictions from sequence and MD data based on linear regression *R*^2^ correlation coefficients, slope and intercept of linear regression curves, and mean squared errors (MSE) between given and predicted phenotypes. Every metric was calculated as the average over 10 sets.

### VAE models of MD data provide structural classification of mutants

Finally, we developed a VAE model based on the MD distance data to identify structural principles underlying different phenotypical outcomes since the sequence-based ML predictor developed above does not provide such insight. The goal of the VAE model was to reduce the high-dimensional intra-molecular distance network into a low-dimensional latent space and then analyze clusters in latent space to extract the key structural determinants giving rise to different phenotypes. The VAE model was applied as described previously (35), but we also added an attention layer both on the encoder and decoder sides. The resulting latent space mapping is shown in Fig. 6A. Mutants in latent space were further grouped into three clusters using a Kmeans clustering algorithm. Although prediction models from MD could not predict phenotypes well, the VAE models did result, to some extent, in a phenotypical separation of mutants as average phenotypes between the clusters I, II and III differed with average phenotypes of −0.49, −0.75 and −0.83, respectively (see Table S2). Cluster I has a mixture of phenotypes with GOF and LOF mutants as the majority, and cluster II and III contains a majority of lethal and LOF and a minority of GOF mutants.

**Figure 6.**
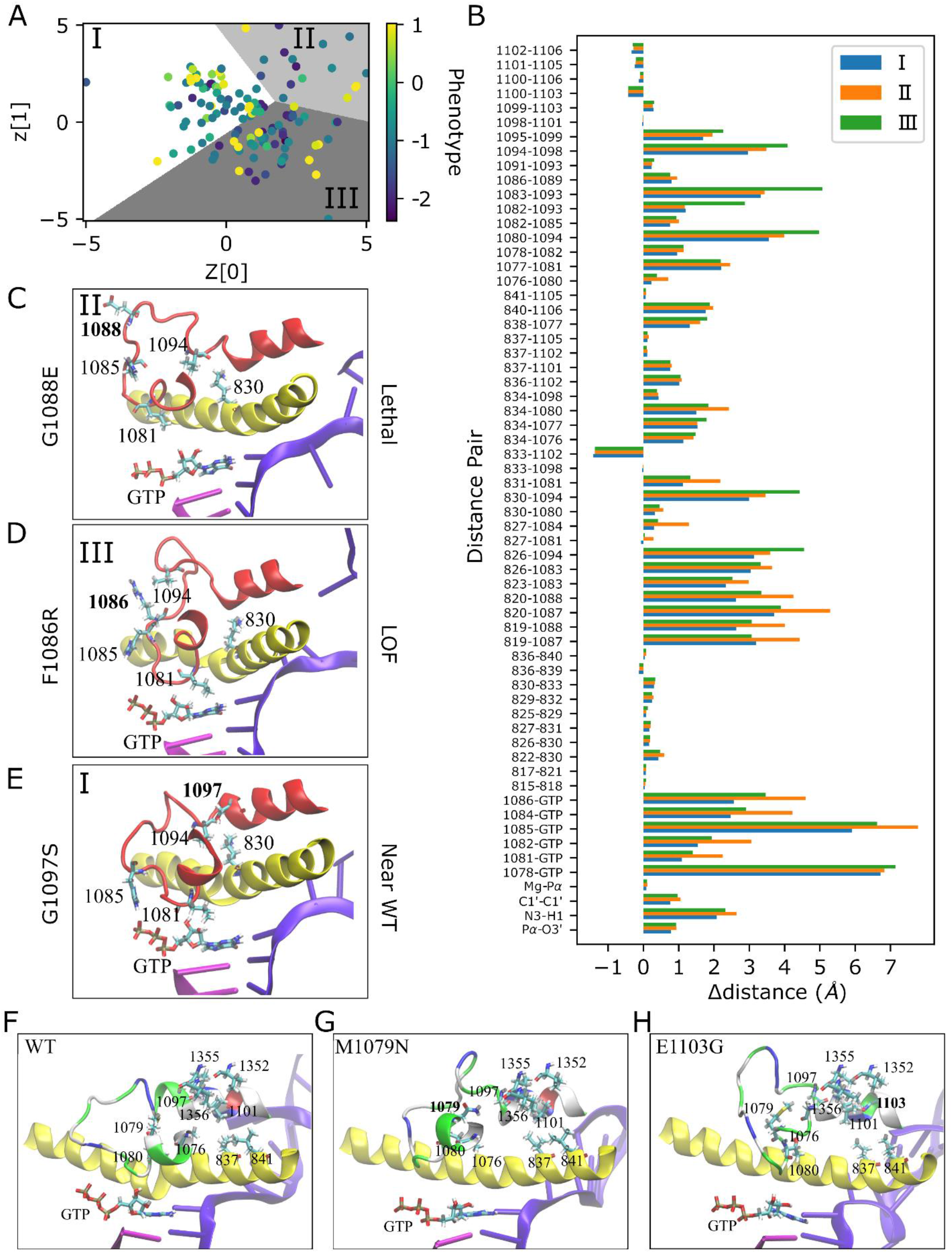
VAE analysis of MD distance data. (A) Latent space from VAE model based on MD distance data shaded according to clusters (I is white, II is gray, III is dark gray) and with each point colored to the mutant phenotype. (B) Distance difference graph; Δdistance shows the differences of the distances from the generative model of VAE at the cluster centers from the WT structure after the equilibration. Panels (C), (D) and (E) show the active site neighborhoods for representative mutants for each cluster in final MD snapshots with the TL (red), BH (yellow), DNA (violet), RNA (magenta) and GTP. Atoms are colored by atom name for residues shown in licorice representation. Panels (F), (G) and (H) show the residues forming hydrophobic pocket for the WT after equilibration and GOF mutants at their final MD snapshots. Color codes are the same as described above except that the TL is shown with residues colored by residue type where positively charged amino acids are in blue, negatively charged ones are in red, polar ones are in green, and hydrophobic ones are in white.

We proceeded to identify cluster centers in latent space and then applied the generative decoder model to the cluster centers to obtain representative distance information for each cluster. Fig. 6B shows the difference map between WT and the resulting distances of each cluster based on the latent space center coordinates. The distances for cluster I, which is dominated by WT-like and GOF mutants, are generally lower than for the other two clusters with more lethal and LOF phenotypes. The finding of longer intra-molecular distances for key residues around the TL for LOF/lethal mutants is similar as to what is shown above from the direct analysis of MD data (Fig. 4C). However, more detailed insights can be extracted from the VAE-based analysis: Cluster II features larger distances between TL residues and GTP compared to the other clusters, especially for H1085 and L1081 suggesting that the underlying mutants (T1077N, R; T1080W; F1084D, N; F1086S; G1088E; S1091W; N1106F) act by directly distorting the active site and thereby hindering catalysis. Cluster III also features larger distances between TL and GTP for some mutants (Q1078C, K, S, W; L1081A; H1085L, Q, S, W; A1087Y, V1089G, P1099D). However, cluster III mostly features increased distances for BH-TL pairs, especially for the distances of the residues K830 and D826 with V1094 that are known to be in close distance in the closed TL (4). This suggests a more indirect mechanism for affecting the mostly negative phenotypes in this cluster that involves disrupting the BH-TL interactions. The BH-TL interactions have been suggested to be functionally important previously (7, 32). In cluster III, most of the substitutions causing large K830-V1094 distances are at close distance to BH, while there are also mutants that are away from BH and results in large distances between K830 and V1094 (see Fig S8). Some of the TL-TL distances for the residues that are close to V1094 are also increased for the cluster III like the distances of T1083-K1093, N1082-K1093, T1080-V1094. This suggests, furthermore, a particularly important role of V1094 for the transcription mechanism. Individual members of each clusters show a similar trend with the cluster centers (Fig. S9); the members of cluster II, including the mutants distant to the GTP that are N1106F, S1091W and G1088E, show larger distances for GTP and trigger loop residues while the members of cluster III (F1086R, Q1078K, L1081Y, A1087K and T1080K given in Fig. S9) have higher TL-BH distances for mutants both close to BH (L1081Y, A1087K and T1080K) or away from BH (F1086R and Q1078K).

To further illustrate structural details, we selected three representative mutants for each cluster based on the following key distances: 1085-GTP, 1081-GTP, 830-1094. Accordingly, we selected the G1088E, F1086R and G1097S with predicted numerical phenotypes of −1.75, −0.92 and 0.33 based on fitness data, for clusters II, III, and I, respectively. G1088E would be considered a lethal mutant (phenotype ≤ −1.5). As a result of the mutation, there are large distances between GTP and 1085 (Fig 6C) which are expected to inhibit enzyme function. F1086R is a LOF mutant where distances between GTP and residues 1085 and 1081 remain relatively close, but the distances between BH and TL increase as seen for the pair of 1094 and 830 (Fig. 6D). As a result, catalysis is still likely possible but less efficient. Finally, G1097S exhibits a near-WT phenotype as it has 0.33 predicted numerical phenotype, which can be classified as weak GOF-near WT. In this mutant, all residues surrounding the active site are close, resulting in the tighter active site geometry (Fig. 6E) that is necessary for WT-like enzyme performance.

GOF mutants are scattered mostly between clusters I and III suggesting that there may also be different mechanisms for this phenotype. The main difference is that the GOF members of cluster III demonstrated relatively larger distances for BH-TL residues compared with GOF mutants in cluster I (Fig. S10). We selected two mutants, M1079N and E1103G in the clusters III and I, respectively, as representative examples. Their final snapshots are shown in Fig. 6F-H with a focus on the hydrophobic pocket formed by the TL residues A1076, M1079, T1080, G1097 and L1101. Both M1079N and E1103G lead to disruptions of the hydrophobic pocket although the mechanism may be slightly different. M1079 points directly to the hydrophobic pocket with close distances to V1355, V1356 and L1101 in the WT equilibrated structure. Mutation of M1079N directly disrupts the hydrophobic interactions. The last snapshot of the E1103G simulation also shows a disrupted hydrophobic pocket with the M1079 pointing away, but because 1103 is positioned further away, the effect must be more indirect by modulating TL-BH interactions. These results support the hypothesis that the primary mechanism for GOF phenotypes is via the direct or indirect disruption of the hydrophobic pocket formed by TL residues near the BH as suggested by previous studies (30, 32).

## DISCUSSION

In this study, we applied ML approaches to interpret genetic fitness values, sequence information, and MD simulation data to predict and characterize TL mutants of yeast RNA Pol II. Fitness values from different conditions were used to generate a quantitative score for TL mutants on a phenotypic continuum from GOF to LOF and lethal. Then, we asked if machine learning approaches using protein sequence and MD simulation could predict these phenotypes when trained on a subset of mutants. The amino acid sequence of proteins is widely used with machine learning approaches to predict various information about structure (36–40) and function (41–43) to mutational phenotypes (44–47). These studies suggest that sequences contain the key information about the structure and function of the proteins, which can be learned. In this study, we used the sequence of TL residues as an input and obtained predictive models of the mutant phenotypes that provide higher correlations than a simple model from average phenotypes for each residue. It may be possible to improve these models further by incorporating additional data on basic physical properties of mutation sites like molecular weight and volume, hydrophobicity, surface area, solvation energy, electrostatic interactions, position specific scoring matrix (PSSM), *etc*. as applied in some of earlier studies (44, 45, 48). Similar approaches could be used for RNA polymerase or other systems to understand function and phenotypes, especially for disease-related mutants. We also tested the sequence-based models trained on single mutants for the prediction of double mutant phenotypes. The model predicts the addition of similar type phenotypes and suppression of opposite type phenotypes and it predicted lethal phenotypes for some LOF-LOF double mutants like H1085Q-Q1078S, but was unable to predict lethal phenotypes from the combination of GOF-GOF mutants observed previously(21). Prediction could easily be improved by training with double mutants allowing the model to learn different double mutant effects.

Recent studies showed that the combination of MD simulations with ML algorithms could provide insight on dynamics, conformations, and kinetics of proteins (49–53). With the motivation from these studies, we used the distance data from MD simulations to investigate their predictive performance. There are computational limitations of working with a large protein like RNA Pol II. Thus, instead of running long simulations, we performed multiple, relatively short simulations to obtain insight about the structural effects of mutations. Surprisingly, we found that structural data obtained from these MD trajectories could not add predictive abilities with respect to the functional phenotypes for specific mutations when used within a ML framework. In light of this finding, several potential advances can be imagined. First, longer, perhaps on μs scales, or higher-quality simulations with different force fields might allow greater inference on mutant mechanisms. Second, obtaining additional data from simulations using a starting structure with an open TL in addition to the closed TL structure may be needed to provide more detailed structural, physics-based input in addition to what is already encoded in differences in amino acid sequences when predicting enzyme function.

Despite the limitation in the MD data, VAE models developed based on MD still could provide mechanistic insights into the potential structural basis of lethal, LOF, and GOF mutants. Based on this analysis, it appears that lethal and more serious LOF phenotypes correlate with mutations that directly increase the distance between key TL residues and the GTP in the active site, thereby inhibiting catalysis. Weaker LOF phenotypes for the examined mutants appear to be related more to disruptions in the TL-BH interaction. In contrast, a substantial fraction of GOF phenotypes appear to result largely from a disruption of a hydrophobic pocket near the TL, and as a consequence closing of the TL is favored, with fidelity presumably reduced. In most of these GOF mutants, we observed increased BH-TL distances especially for the cases when the mutants located close to BH. These general findings are consistent with previous studies but go further because of a more comprehensive analysis that is based on a systematic analysis of a larger number of TL mutations. Still, more work is left to be done to understand specific mutations and specific roles of individual residues. One avenue for further studies may be via the exploration of double mutants that may restore non-WT phenotypes to WT function based on the predictions made above.

## CONCLUSION

In this study, we report a comprehensive characterization of the mutations of RNA Pol II TL residues using ML techniques on high throughput genetic fitness data, sequence data and MD simulations of TL residue mutations. Our study suggests that fitness data and sequence information are correlated such that the phenotype that was predicted from fitness values can be learned by sequence information and the predictions from sequence go beyond the simple prediction model from average phenotypes. Such a prediction was not possible with our MD data due to the computational limitations. Nevertheless, the MD data could provide some mechanistic understanding on different phenotypes. Longer simulations may be necessary to obtain a predictive model that would be comparable with the sequence-based model. However, μs scale simulations of large TL mutation libraries of RNA Pol II still remain a major computational challenge. As an alternative to that, artificial intelligence methods that predict structural details from sequence of mutants could be developed and applied to RNA Pol II systems as a future direction.

## METHODS

### Prediction of Continuous Phenotypes from Fitness Data

The Pol II TL fitness and phenotypic landscape approach of Qiu *et al*. based on a deep mutational scanning approach was applied here to a second-generation TL mutant library. The construction of the TL mutant library was followed by *en masse* phenotyping assays monitored by deep sequencing. This work will be described in detail elsewhere (see supplied Methods for review) but key updates to the original approach are as follows. A second-generation TL mutant library was synthesized using programmed oligo synthesis (Agilent). Library oligos were amplified from synthesized pools and homology arms of WT sequence were added using overlap PCR. These fragments comprising ∼200 nt of flanking *RPB1* sequence on each side and a central 93-nt region encoding the WT Pol II TL (Rpb1 amino acids 1076-1106) or individual TL variants were introduced into yeast along with *RPB1* plasmid lacking TL sequence and linearized at the TL position in three replicates. This cotransformation allows construction of a pool of variant plasmids by gap repair, exactly as performed previously by Qiu et al. Transformants were plated at high density (∼10,000 per plate) instead of 300-400 as done previously. 5-FOA-resistant colonies were scraped from SC-Leu+5FOA plate and replated on SC-Leu, SC-Leu + 20mg/ml MPA (Fisher Scientific), SC-Leu + 15 mM Mn (Sigma), YPRaff, YPRaffGal, SC-Lys, and SC-Leu + 3% Formamide (JT Baker) plates for phenotyping. Cells were scraped from each phenotyping plate after defined growth periods and genomic DNA was extracted from each screening plate with Yeastar Genomic DNA kit (Zymo) and amplified using emulsion PCR (EURx Micellula DNA Emulsion & Purification (ePCR) PCR kit) according to manufacturer’s instructions. A dual indexing strategy where custom indexing primers were paired with primers using 28 NEB indices was utilized in the amplification to discriminate between various screening plates. Amplified libraries were sequenced with Illumina Next-seq for 150nt single-end reads.

Experimental fitness data for yeast Pol II TL variants from a total of 21 conditions with three replicates each were used for deriving a predictive phenotype model. Missing fitness values were imputed using mean values of a given feature. The model was based on continuous real-valued phenotypes, where previously classified GOF, LOF, and lethal outcomes map to values of +1.0, −1.0, and −2.0 and where the WT maps to 0.0. A neural network consisting of three fully-connected layers with 256, 128 and 64 nodes (see Fig. S11A) was trained against known phenotypes from previous studies (3, 21, 32) for 83 mutants. Mutations that did not lead to clear GOF, LOF, or lethal outcomes were treated as WT with a continuous phenotype value of 0.0. The mean squared error (MSE) was used as the loss function. After training based on the classified mutations, phenotypes were predicted from the experimental fitness data for all of the TL mutants.

The fitness data was used further in a variational autoencoder model to map the phenotypes into a reduced dimensional space. Both encoder and decoder models have three layers with 256, 128 and 64 nodes (see Fig. S11B). The loss function of MSE between input features and generated output values was used. For the regularization of the latent space, the Kullback-Leibler (KL) divergence between latent space distribution and the standard normal distribution was applied as defined by Kingma and Welling (35). All the fitness data, the predicted phenotypes, and the latent space coordinates are summarized in the Supplementary Spreadsheet 1.

### MD Simulations

The WT RNA Pol II structure used as a starting point was deposited to Protein Data Bank with PDB ID:2E2H(4). Missing residues were modeled for the loops that have less than eight amino acids using MODELLER version 9.15 (54). The histidine at 1085 of Rbp1 was protonated based on the study by Huang et al (18). The system was solvated in a cubic box with a 10 Å cutoff distance between the box edges and any atom of the RNA-Pol II complex resulting in a total box size of 162 Å. The system was then neutralized with Na^+^ ions. Periodic boundary conditions were applied along with the particle-mesh Ewald Algorithm for the calculation of long-range electrostatic interactions. Lennard-Jones interactions were switched from 10 to 12 Å. The SHAKE algorithm was used to constrain bond lengths involving hydrogen atoms. The CHARMM 36m force field (55) was used for proteins and the CHARMM 36 force field (56) was used for nucleic acids. The TIP3P model (57) was used for explicit water molecules and a recently suggested NBFIX for Na^+^ phosphate interactions was applied (58). The force fields were modified to redistribute the atomic masses of atoms that are attached to hydrogen atoms, so that hydrogen atoms had an increased mass of 3 a.m.u instead of 1(59). This modification allowed us to perform the simulations using a 4 fs time step.

The WT system was subjected to 5,000 steps of energy minimization. The system was then equilibrated for around 1.6 ns by gradually increasing the temperature from 100 K to 300 K and using restraints on the heavy atoms of backbone and sidechains with force constants of 400 and 40 kJ/mol/nm^2^, respectively. The equilibrated Pol II complex was used to prepare the single site TL mutants. In total, 135 mutants were prepared (see Supplementary Spreadsheet 2). Each mutant system was minimized again using the same minimization scheme as for the WT and equilibrated for an additional 1 ns. Production simulations were then performed using Langevin dynamics with a friction coefficient of 0.01 ps^-1^ under the constant temperature of 298 K. Simulations were run using OPENMM(60) on GPU hardware. 100 ns production runs were performed for three replicates for each mutant and the last 50 ns of the production runs was used for the analysis. In total, 40.5 μs of mutant simulations were generated.

The analysis of the MD simulations was done using the MMTSB package(61) in combination with in-house scripts. The minimum distances between the residues were analyzed and the average distances were calculated from the combined distribution of distances from the three replicate simulations. The RMSD was calculated for the TL and BH regions for all the mutants and is shown in Figs. S12 and S13, respectively.

### Phenotype prediction models using sequence and MD data

For models predicting phenotypes based on the TL amino acid sequence, sequence data was converted into one-hot encoding sparse matrices, where each feature was a 21-size vector that represents a single amino acid along the sequence. Histidine was coded as either protonated or deprotonated to allow the model to distinguish protonated histidine at residue 1085. The one-hot-encoded sequence-based features, a two-dimensional matrix (31×21), were used as input for a neural network with three fully connected layers with 128, 64, and 32 nodes in two dimensional matrices (31×128, 31×64, 31×32), respectively. The 32-node third layer was flattened two a one-dimensional vector with a size of 992 and passed through another layer with 32 nodes before the single-valued output layer (see Fig. S11A).

For models trained to predict phenotypes based on MD data, pairwise distances between key residues involving the TL and neighboring elements were extracted from the simulation trajectories and averaged. A list of distances used in the models is provided in the Supplementary Spreadsheet 2. A neural network with three fully connected layers with 128, 64 and 32 nodes was used as for the sequence-based predictors. To emphasize the effect of mutations at the different parts of the structure, we also tested another model that has 128 and 64-nodes layers that were connected to the output layer via an attention layer (62) (see Fig. S11A). The attention layer was generated for the last hidden layer with 64 nodes, which was used as query, key, and value vectors for the self-attention framework. The new values were calculated by the multiplication of the values and the weighted sum of the similarities between the query and key vectors.

The mutants examined via MD simulations were used for both sequence and MD prediction models to compare the performances of the neural network models. The mutants were split into ten training and test sets with randomly chosen 100 and 35 mutants, respectively. A 1:1 combination of loss functions of mean squared error (MSE) and KL divergence was applied to the prediction models with sequence and MD data. The KL divergence was calculated analytically as follows:

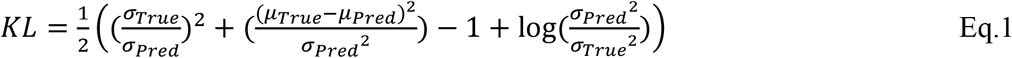

The model weights and biases were saved every 500 epochs and training continued until a maximum of 20,000 epochs. For each set, ten different models were generated, and the best model was chosen from the saved models at intervals that provides the best combination of *R*^2^ correlations and slope for the regression line between predictions and label phenotypes for the test set. The overall performance of the models was calculated by the average performance of the best models of the sets for the comparison of different models from either sequence or MD data. A complete phenotypical landscape was generated using the sequence-based models based on the ensemble average of the predictions from the best models of the sets.

The models based on MD data and sequence were also combined to see if there was any improvement in the predictions when using both data as input features. The data was combined in two different ways: First, we took the pre-trained sequence model and added the MD-based model to be trained to add additional input to the last layer of the sequence model before generating the output. Second, we used a pre-trained MD model and added the sequence model and concatenated the last layers of the two models to generate the final output. In each case, we froze the pre-trained weights to test whether the additional input features can improve the predictions.

### Variational Auto-Encoder Models based on the MD data

Variational auto-encoder models were generated based on the MD data in order to extract mechanistic insights from the simulations. Both, the encoder and decoder networks, consisted of three fully-connected layers with 128, 64, and 32 nodes. We applied an asymmetrical VAE model at which the attention layers were applied after the layers with 32 nodes for both, the encoder and decoder (see Fig. S11B). We experimented with 2D and 3D latent spaces. The performance of the generative model with a 3D latent space was similar to the model with the 2D latent space (see Fig S14). Therefore, we performed further analysis of the model with the 2D latent space. The resulting reduced dimensionality latent space was then clustered via a Kmeans clustering algorithm. Three clusters provided the best separation of average phenotypes between the clusters (see Fig S15). Thus, we separated the latent space into three clusters. The cluster centroids were used subsequently to generate representative molecular states for each cluster via the decoder network.

### Machine learning details and software

All ML models were generated with the Tensorflow package (63). The models are summarized as diagrams in Fig S11. The Python scripts to train the models and to predict from the trained models along with the weight of the trained models and the input files for the models are available at https://github.com/bercemd/PolII-mutants. Data imputation, Kmeans clustering and principal component analysis (PCA) were performed with the Sklearn module in Python (64). The Adam optimizer was used for all models. The learning rate and number of epochs were varied according to the model that are summarized in Table S1. Different learning rates were applied to prevent unstable training but achieve convergence to a minimum loss within reasonable training times (Figs. S16-18). A batch size of 4 was used for all models except for the prediction models with the KL divergence loss in which a batch size of 100 is used to calculate the loss for the complete training set. The rectified linear unit (ReLU) activation function was used for each hidden layer for both prediction and VAE models except for the attention layer, where the Softmax activation function was used.

## ACKNOWLEDGMENTS

We thank Guillermo Calero for helpful discussions. This study was funded by the National Institutes of Health (R35 GM126948, to MF, R01 GM097260 and R35 GM144116 to CDK). We used the computational resources at the Institute for Cyber-Enabled Research/High Performance Computing Cluster (ICER/HPCC) at Michigan State University and at the National Science Foundation’s Extreme Science and Engineering Discovery Environment (XSEDE) facilities under the grant TG-MCB090003.

## AUTHOR CONTRIBUTIONS

**Bercem Dutagaci**, Conceptualization, Data Curation, Formal Analysis, Investigation, Methodology, Project Administration, Software, Validation, Visualization, Writing – Original Draft Preparation, Writing – Review & Editing; **Bingbing Duan**, Data Curation, Formal Analysis, Investigation, Methodology, Validation, Writing – Review & Editing; **Chenxi Qiu**, Data Curation, Formal Analysis, Investigation, Methodology, Validation; **Craig D. Kaplan**, Conceptualization, Data Curation, Formal Analysis, Funding Acquisition, Investigation, Methodology, Project Administration, Resources, Supervision, Validation, Writing – Original Draft Preparation, Writing – Review & Editing; **Michael Feig**, Conceptualization, Data Curation, Formal Analysis, Funding Acquisition, Investigation, Methodology, Project Administration, Resources, Software, Supervision, Validation, Visualization, Writing – Original Draft Preparation, Writing – Review & Editing

## SUPPORTING INFORMATION

**Figure S1**. Principal component analysis (PCA) on the fitness data with each data point colored according to its corresponding phenotype.

**Figure S2**. Training and test loss during the training of the models. The models are trained with the input features from sequence data (Sequence), MD data without an attention layer (MD wo attention), MD data with an attention layer (MD w attention), MD and sequence data with pre-trained sequence weights (MD pre-trained sequence) and pre-trained MD weights (Sequence pre-trained MD).

**Figure S3**. Correlations between predictions and label phenotypes for the training sets of different models. The model details are as in Fig. S2.

**Figure S4**. Correlations between predictions and label phenotypes for the test sets of different models. The model details are as in Fig. S2.

**Figure S5**. The difference map for the phenotypes predicted from sequence and fitness. Y-axis shows the absolute value of the differences of phenotypes from the fitness and sequence data and X-axis shows the phenotypes from the fitness data. The outliers with difference larger than 1.25 are shown.

**Figure S6**. Prediction of phenotypes from the average phenotypes for each mutant for the training sets. (A) the phenotypes predicted from the average values vs phenotypes from the fitness and the linear regression line (B) the average phenotypes from the training sets shown in the complete mutation map.

**Figure S7**. Heatmap plot of phenotypes vs average distances between TL residues and GTP (left) and between TL and BH residues (right) for the mutants from MD simulations.

**Figure S8**. The average distances between K830 and V1094 in mutant simulations vs. the minimum distance of the mutated amino acid site to the BH in the WT structure. Each panel shows the plot for the members of each cluster found in the MD-VAE latent space. The dashed lines show the distance of K830 and V1094 in the WT structure.

**Figure S9**. Distribution of distances between L1081 and GTP, H1085 and GTP, and K830 and V1094 for the selected members of the clusters from the VAE latent space. The mutants are given in the legends with their continuum phenotypes in the parentheses.

**Figure S10**. Distribution of distances between D826 and V1094, K830 and V1094, and G819 and G1088 for the selected GOF mutants of the clusters from the VAE latent space. The mutants are given in the legends with their continuum phenotypes in the parentheses.

**Figure S11**. The diagram of the neural network models. (A) The models for the prediction of continuous phenotypes have alternating layers depending on the input: For the models with fitness score as the input, three dense layers were used. For the models with the MD data as the input, three dense layers or two dense layers and one attention layer were used. For the models with amino acid sequence as the input, two-dimensional matrix at the third dense layer was flattened out and passed through another dense layer. (B) VAE model was applied to the fitness scores as three dense layers on the encoder and decoder models. It was applied to the MD data with additional attention layers on the encoder and decoder.

**Figure S12**. RMSD values of TL residues for 135 mutants. RMSD values from three replicate simulations were represented with different colors. There are not large changes in RMSD for TL suggesting that TL is retaining its overall conformation for the mutants within the simulation time scale.

**Figure S13**. RMSD values of BH residues for 135 mutants. RMSD values from three replicate simulations were represented with different colors. There are not large changes in RMSD for BH suggesting that BH is retaining its overall conformation for the mutants within the simulation time scale.

**Figure S14**. The distribution of mutants on the latent spaces (top) and generative performances (bottom) of the VAE models with 3D (left) and 2D (right) latent spaces.

**Figure S15**. The clustering of the VAE latent space of the MD data using Kmeans clustering algorithm with three, four and five clusters. Each cluster is shown in colors from white to different shades of grey; mutants are scattered, and color coded with corresponding phenotypes; at each cluster the average phenotypes of the mutants are shown.

**Figure S16**. Three replicates of VAE models using the fitness data as features at different learning rates. Models with 10^−6^ learning rate were not converged. Models with 10^−3^ learning rate tend to be stuck in a local minimum loss. The models with 10^−4^ and 10^−5^ learning rates provided similar latent spaces without any convergence problem, therefore a learning rate of 10^−4^ was used for fitness-based models.

**Figure S17**. Three replicates of VAE models using the MD data as features at different learning rates. The same trend with the Fig S16 is observed. Learning rate of 10^−4^ provided the most visual separation of phenotypes, therefore it was used for the MD data models.

**Figure S18**. Three replicates of prediction models using the sequence data as features at different learning rates. Learning rate of 10^−5^ provided the minimum losses for the test sets, therefore it was used for the sequence models.

**Table S1**. The summary of deep learning models

**Table S2**. Statistical analysis of the clusters obtained from VAE model using MD data. T-test was performed by assuming the two populations have different variance. Clusters I, II and III corresponds to the clusters shown in Figure 6A

**Supplemental Spreadsheet 1**. The fitness scores, the predicted phenotypes, and the latent space coordinates.

**Supplemental Spreadsheet 2**. The distance analysis results, and the latent space coordinates from the distance input features from MD simulations.

## Notes

https://github.com/bercemd/PolII-mutants

### Competing Interest Statement

The authors have declared no competing interest.

