## Supplementary figures and tables for "Characterization of RNA Polymerase II Trigger Loop Mutations using Molecular Dynamics Simulations and Machine Learning"

**This PDF file includes:**

Figures S1 to S18
Tables S1 and S2

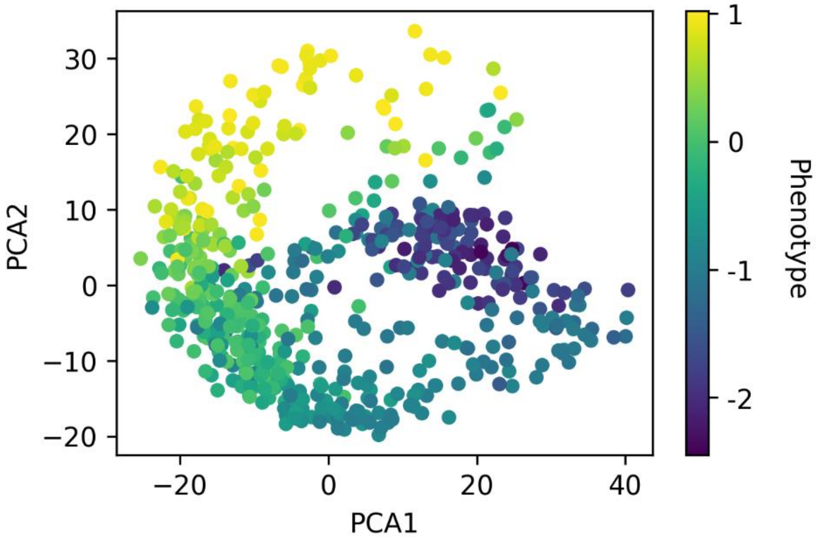


**Figure S1.** Principal component analysis (PCA) on the fitness data with each data point colored according to its corresponding phenotype.

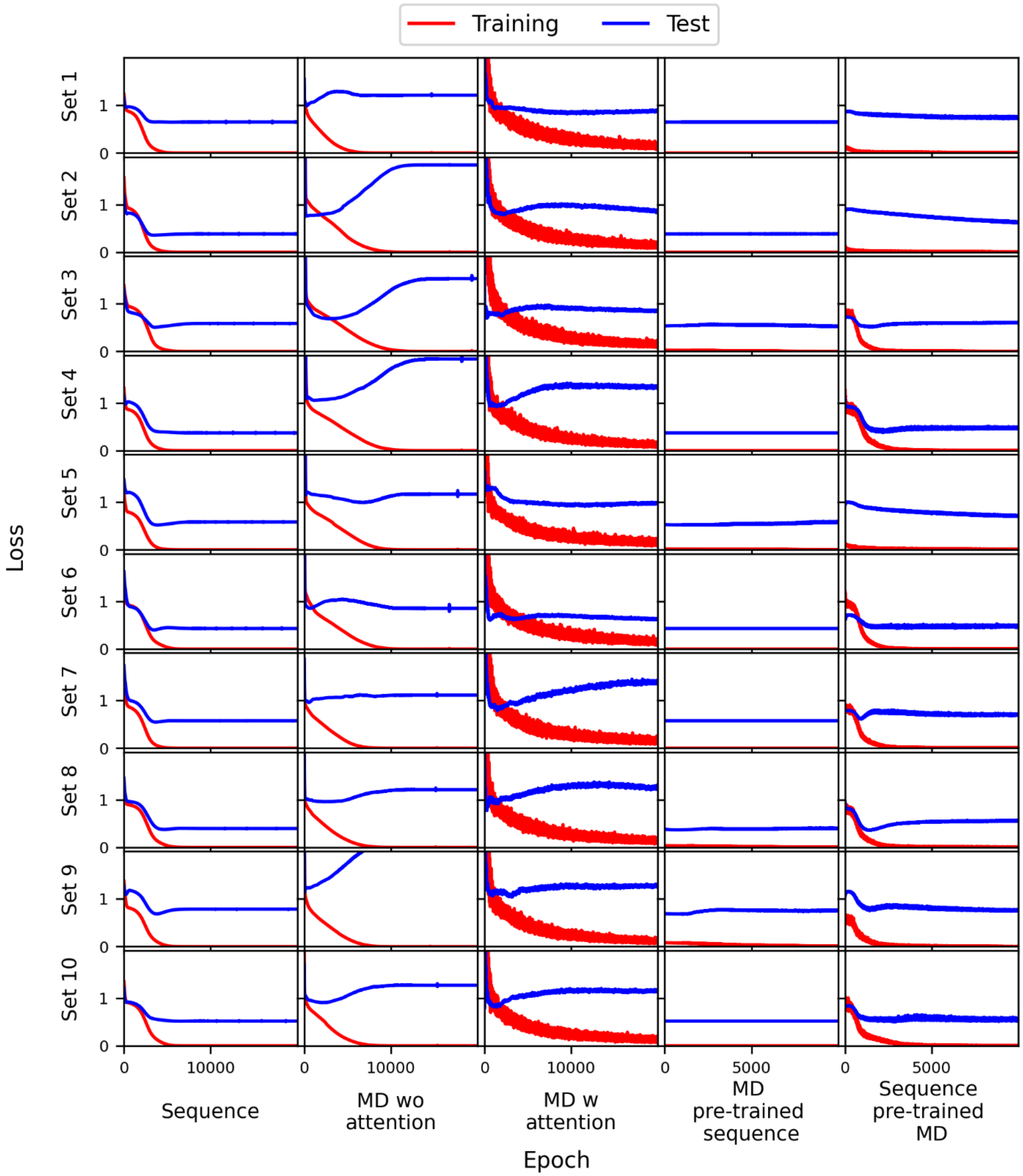


**Figure S2.** Training and test loss during the training of the models. The models are trained with the input features from sequence data (Sequence), MD data without an attention layer (MD wo attention), MD data with an attention layer (MD w attention), MD and sequence data with pre-trained sequence weights (MD pre-trained sequence) and pre-trained MD weights (Sequence pre-trained MD).

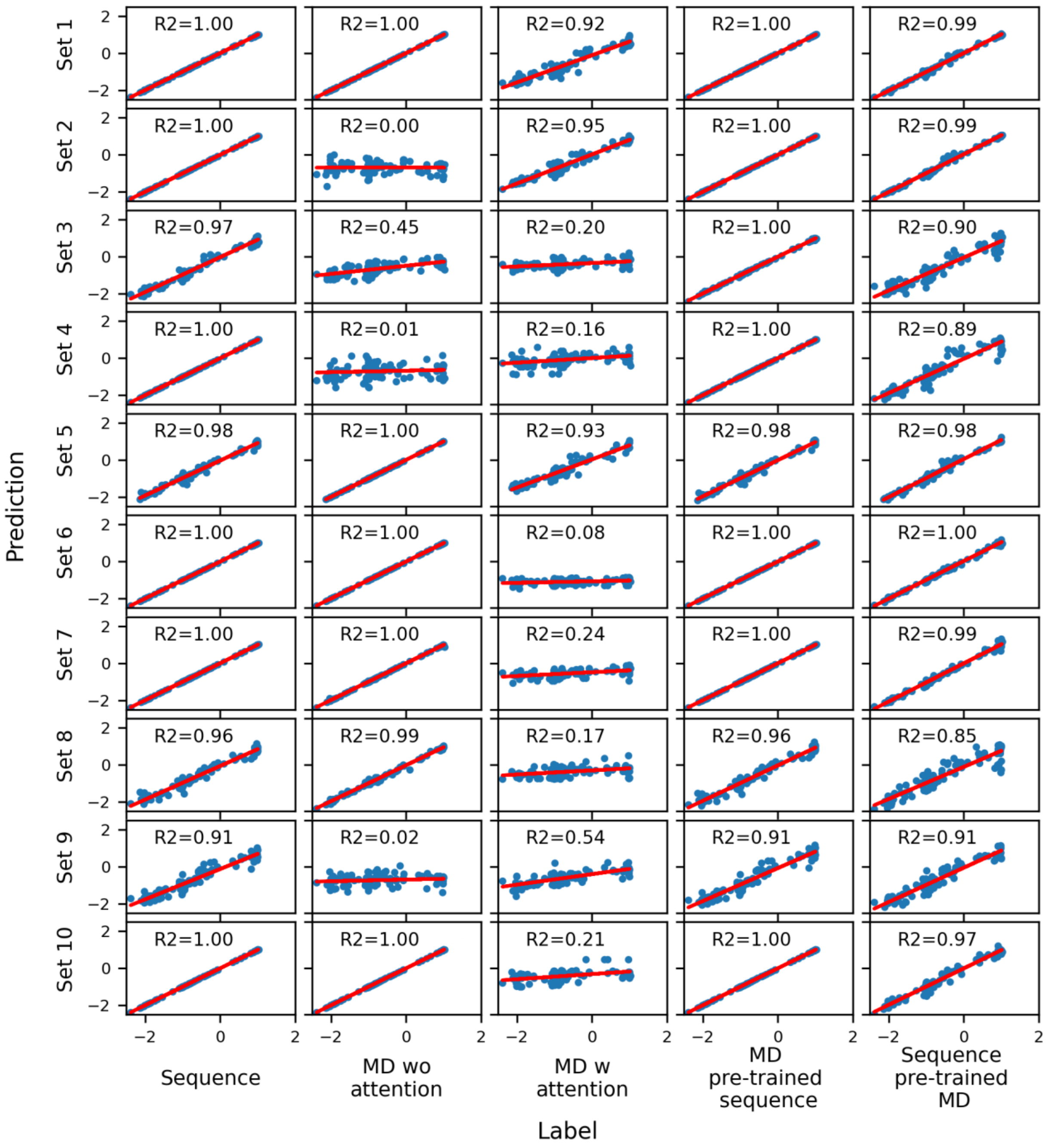


**Figure S3.** Correlations between predictions and label phenotypes for the training sets of different models. The model details are as in Fig. S2.

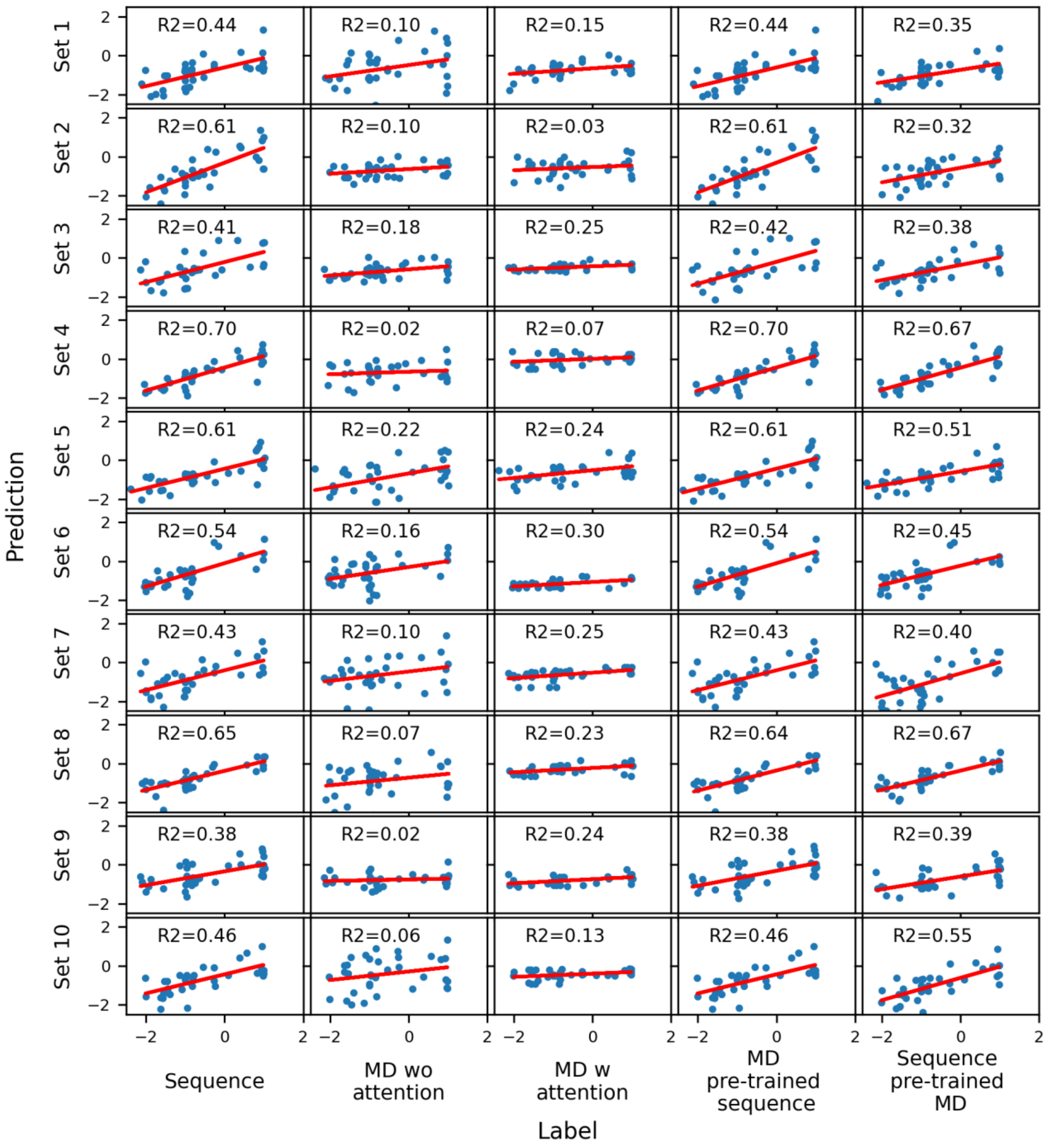


**Figure S4.** Correlations between predictions and label phenotypes for the test sets of different models. The model details are as in Fig. S2.

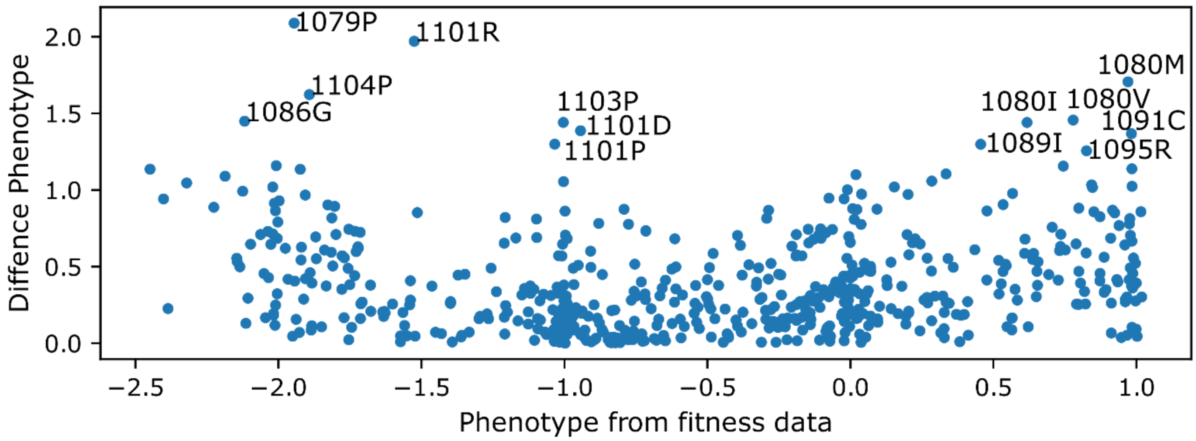


**Figure S5.** The difference map for the phenotypes predicted from sequence and fitness. Y-axis shows the absolute value of the differences of phenotypes from the fitness and sequence data and X-axis shows the phenotypes from the fitness data. The outliers with difference larger than 1.25 are shown.

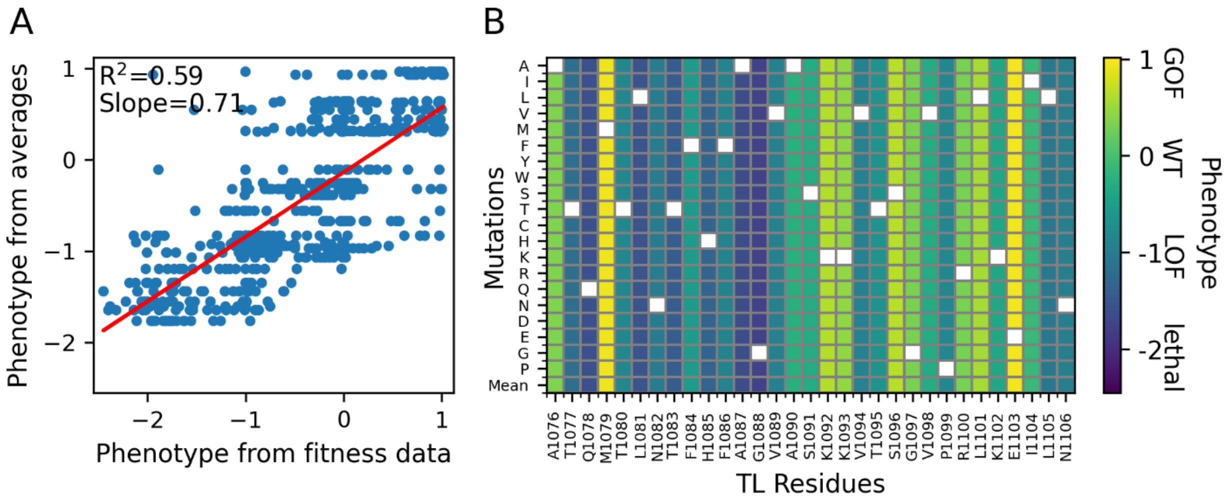


**Figure S6.** Prediction of phenotypes from the average phenotypes for each mutant for the training sets. (A) the phenotypes predicted from the average values vs phenotypes from the fitness and the linear regression line (B) the average phenotypes from the training sets shown in the complete mutation map.

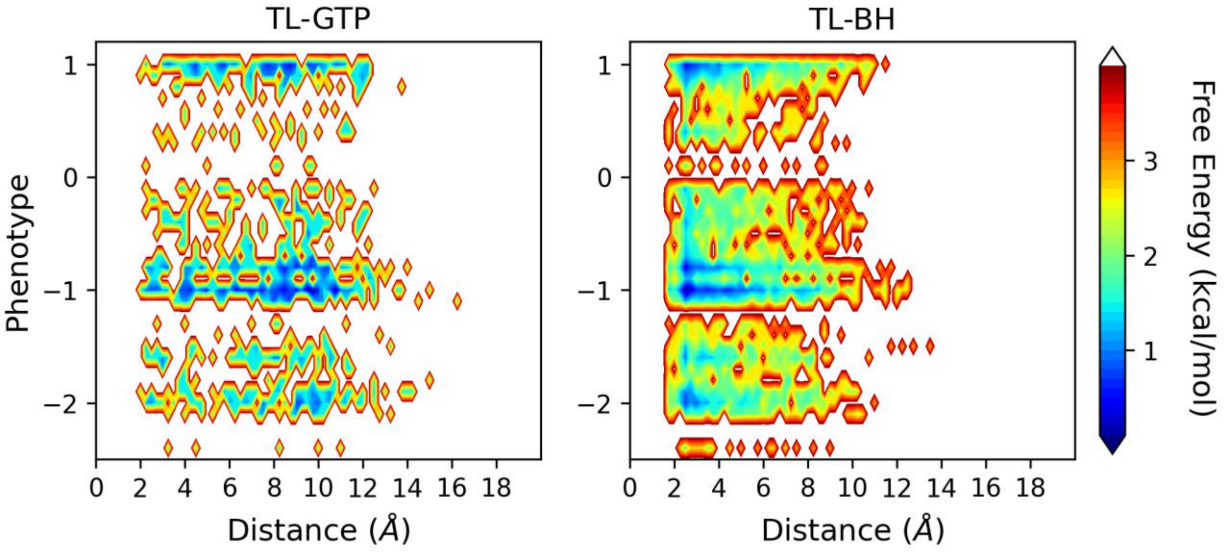


**Figure S7.** Heatmap plot of phenotypes vs average distances between TL residues and GTP (left) and between TL and BH residues (right) for the mutants from MD simulations.

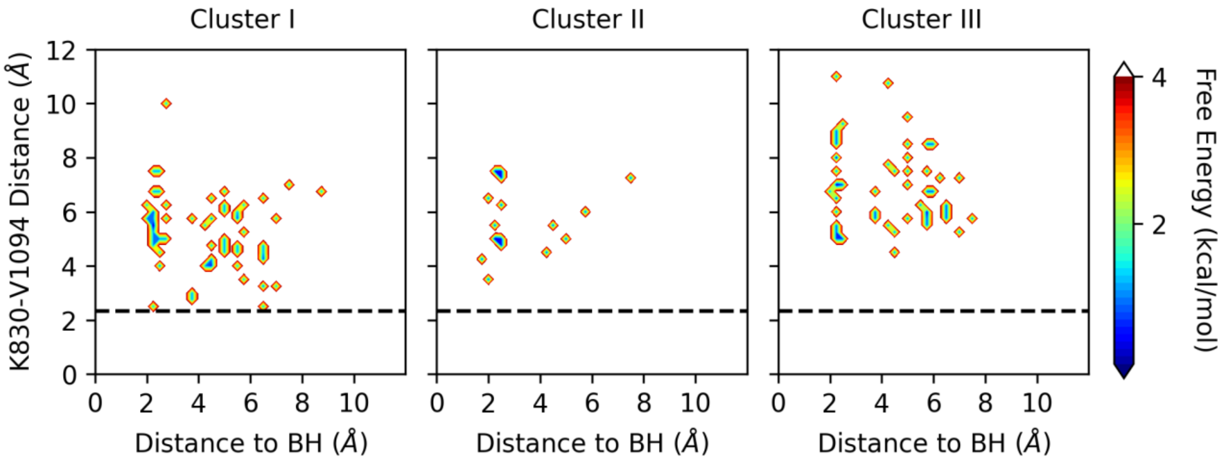


**Figure S8.** The average distances between K830 and V1094 in mutant simulations vs. the minimum distance of the mutated amino acid site to the BH in the WT structure. Each panel shows the plot for the members of each cluster found in the MD-VAE latent space. The dashed lines show the distance of K830 and V1094 in the WT structure.

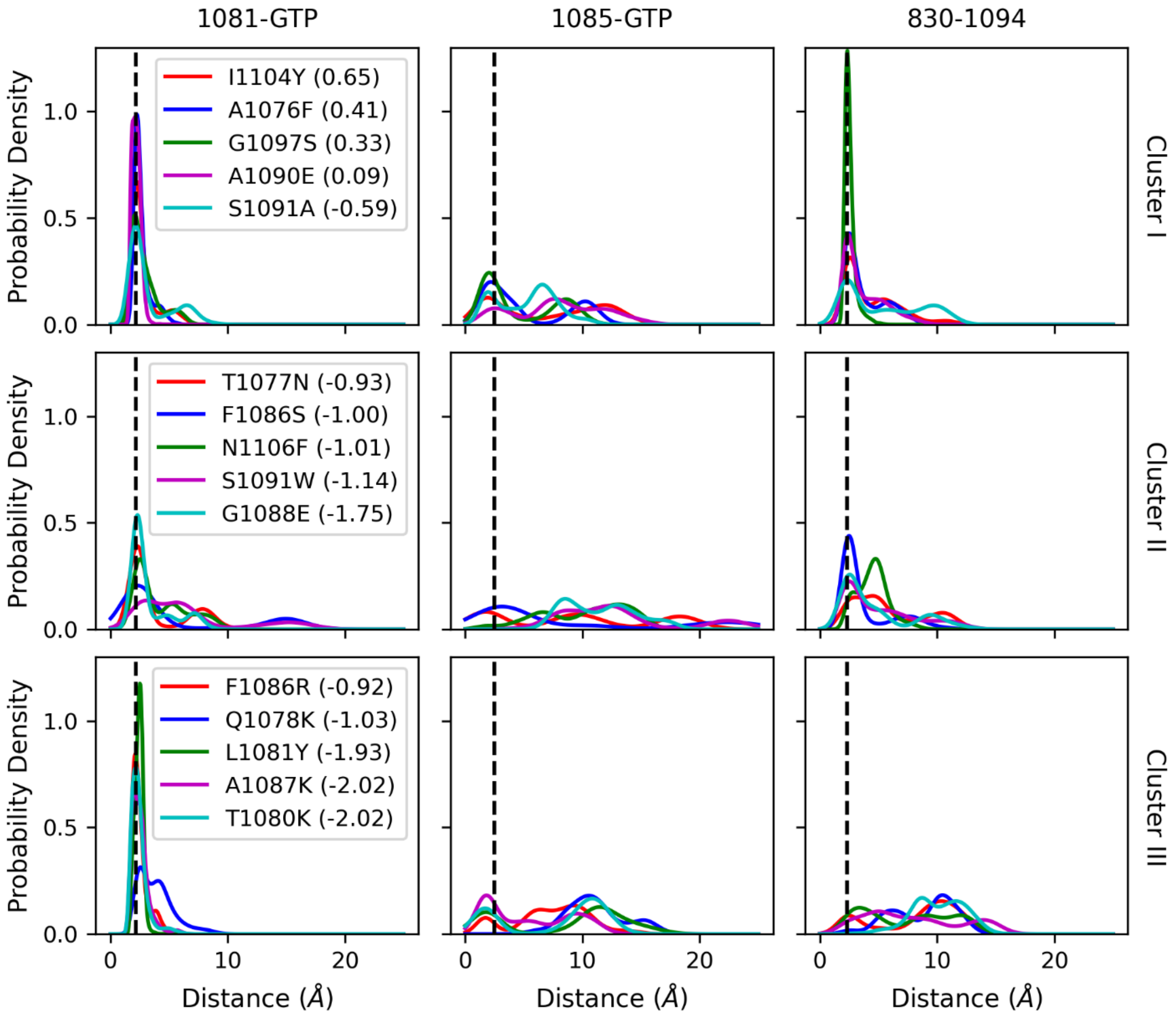


**Figure S9.** Distribution of distances between L1081 and GTP, H1085 and GTP, and K830 and V1094 for the selected members of the clusters from the VAE latent space. The mutants are given in the legends with their continuum phenotypes in the parentheses.

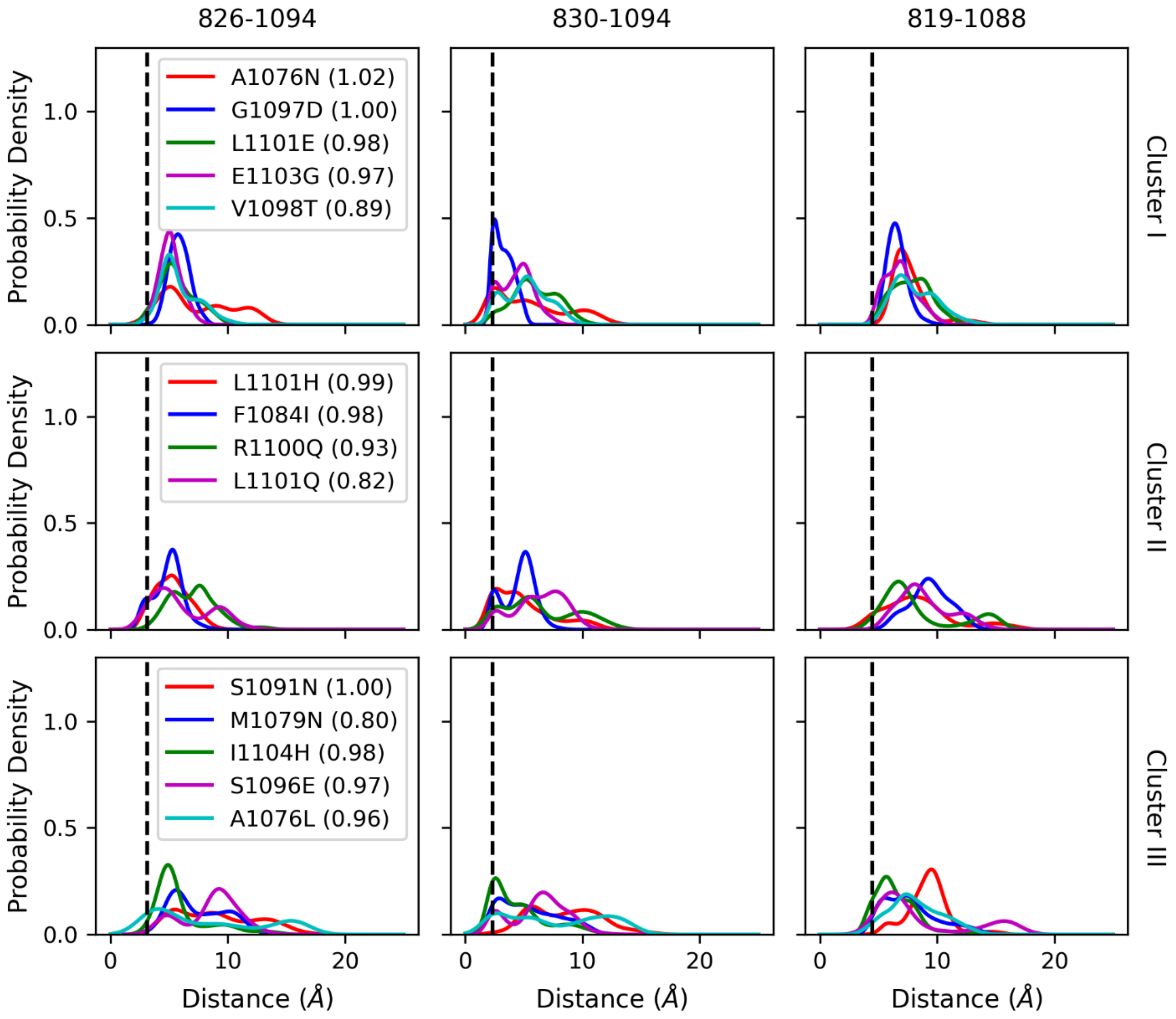


**Figure S10.** Distribution of distances between D826 and V1094, K830 and V1094, and G819 and G1088 for the selected GOF mutants of the clusters from the VAE latent space. The mutants are given in the legends with their continuum phenotypes in the parentheses.

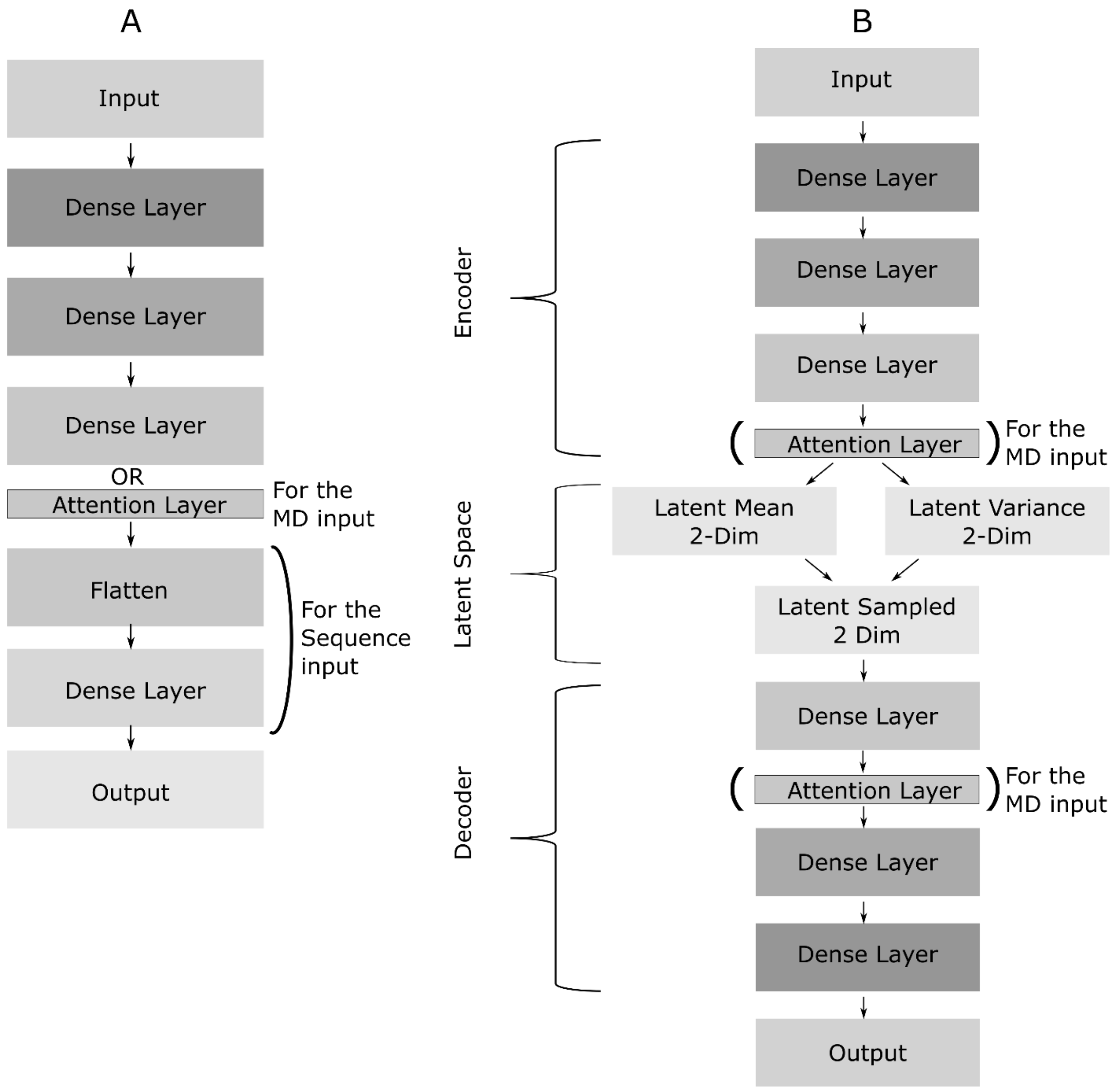


**Figure S11.** The diagram of the neural network models. (A) The models for the prediction of continuous phenotypes have alternating layers depending on the input: For the models with fitness score as the input, three dense layers were used. For the models with the MD data as the input, three dense layers or two dense layers and one attention layer were used. For the models with amino acid sequence as the input, two-dimensional matrix at the third dense layer was flattened out and passed through another dense layer. (B) VAE model was applied to the fitness scores as three dense layers on the encoder and decoder models. It was applied to the MD data with additional attention layers on the encoder and decoder.

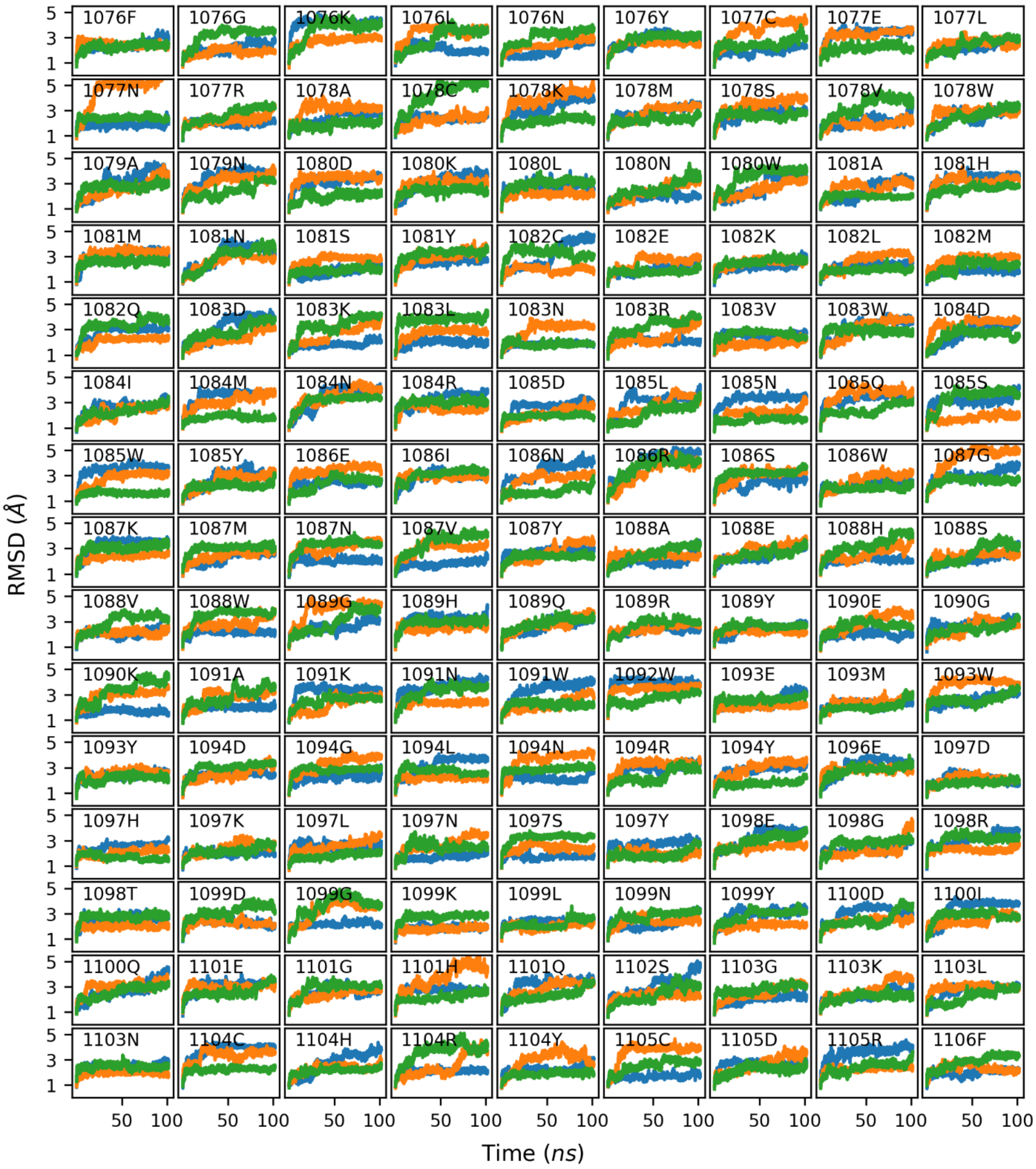


**Figure S12.** RMSD values of TL residues for 135 mutants. RMSD values from three replicate simulations were represented with different colors. There are not large changes in RMSD for TL suggesting that TL is retaining its overall conformation for the mutants within the simulation time scale.

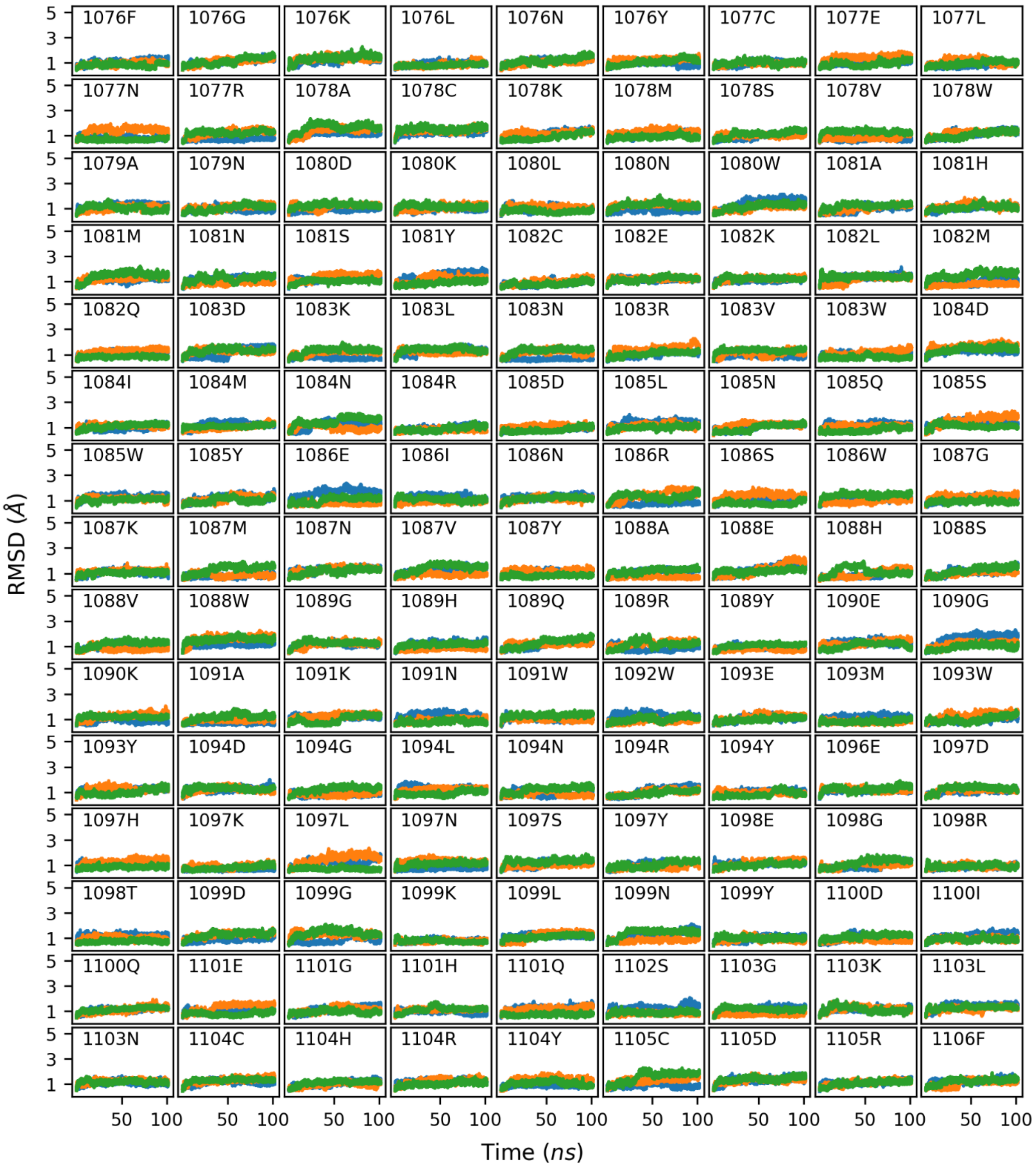


**Figure S13.** RMSD values of BH residues for 135 mutants. RMSD values from three replicate simulations were represented with different colors. There are not large changes in RMSD for BH suggesting that BH is retaining its overall conformation for the mutants within the simulation time scale.

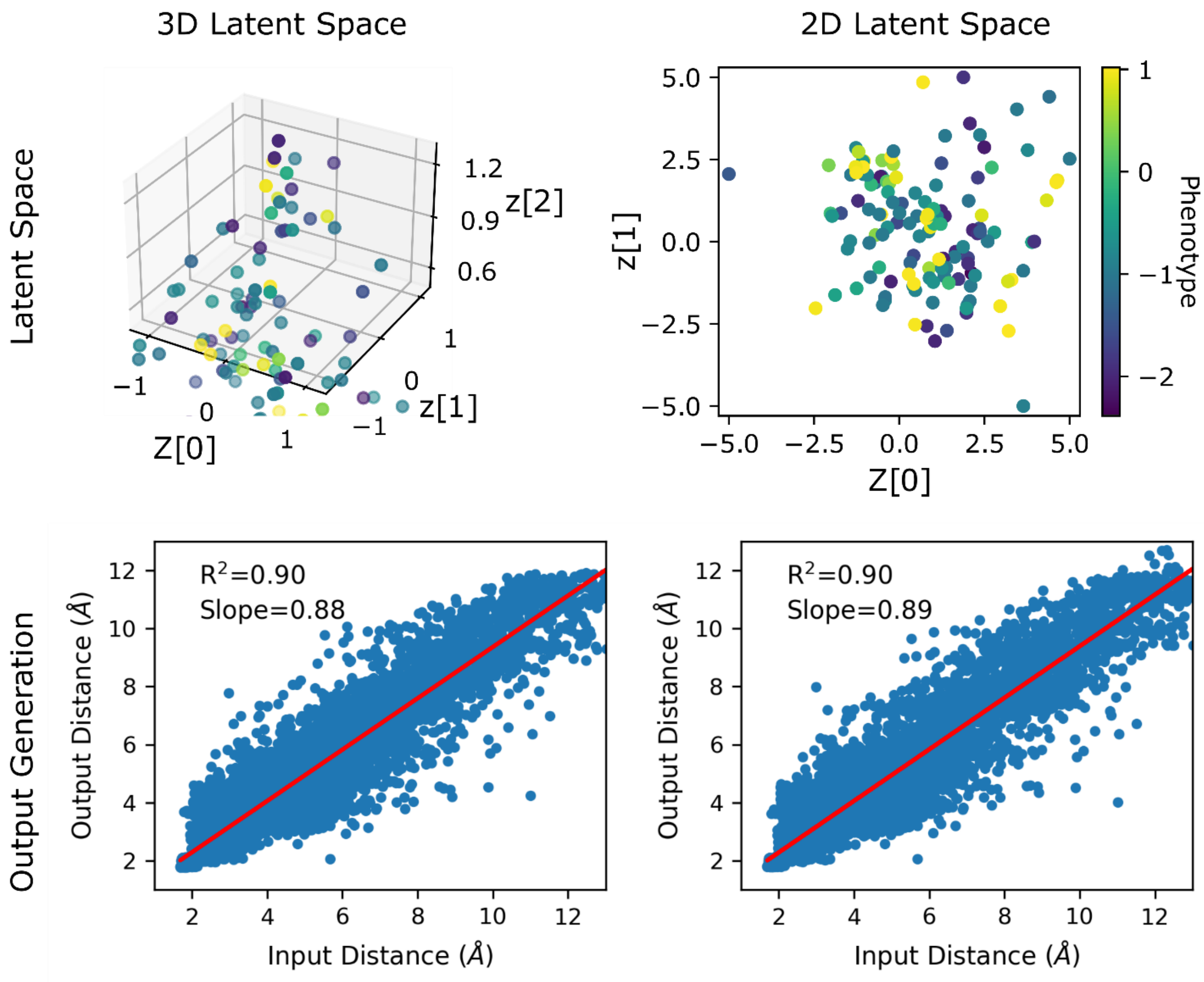


**Figure S14.** The distribution of mutants on the latent spaces (top) and generative performances (bottom) of the VAE models with 3D (left) and 2D (right) latent spaces.

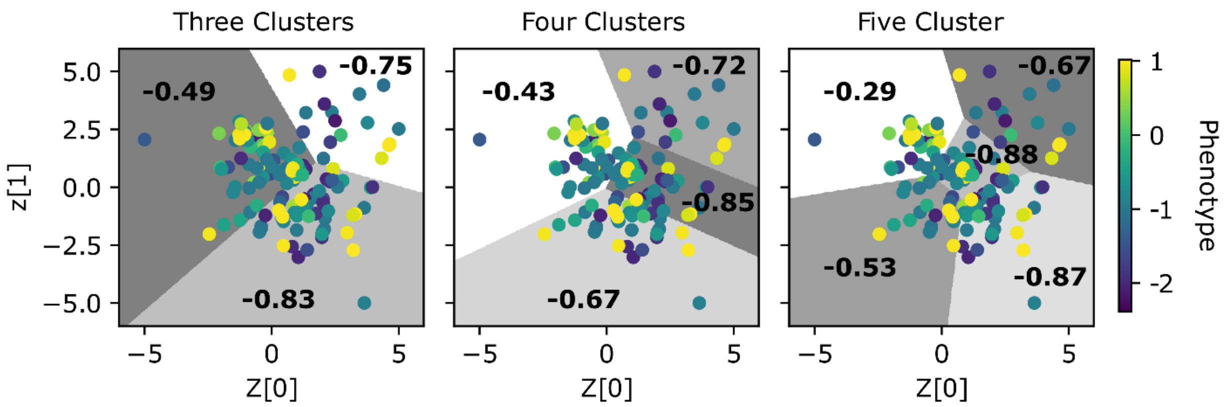


**Figure S15.** The clustering of the VAE latent space of the MD data using Kmeans clustering algorithm with three, four and five clusters. Each cluster is shown in colors from white to different shades of grey; mutants are scattered, and color coded with corresponding phenotypes; at each cluster the average phenotypes of the mutants are shown.

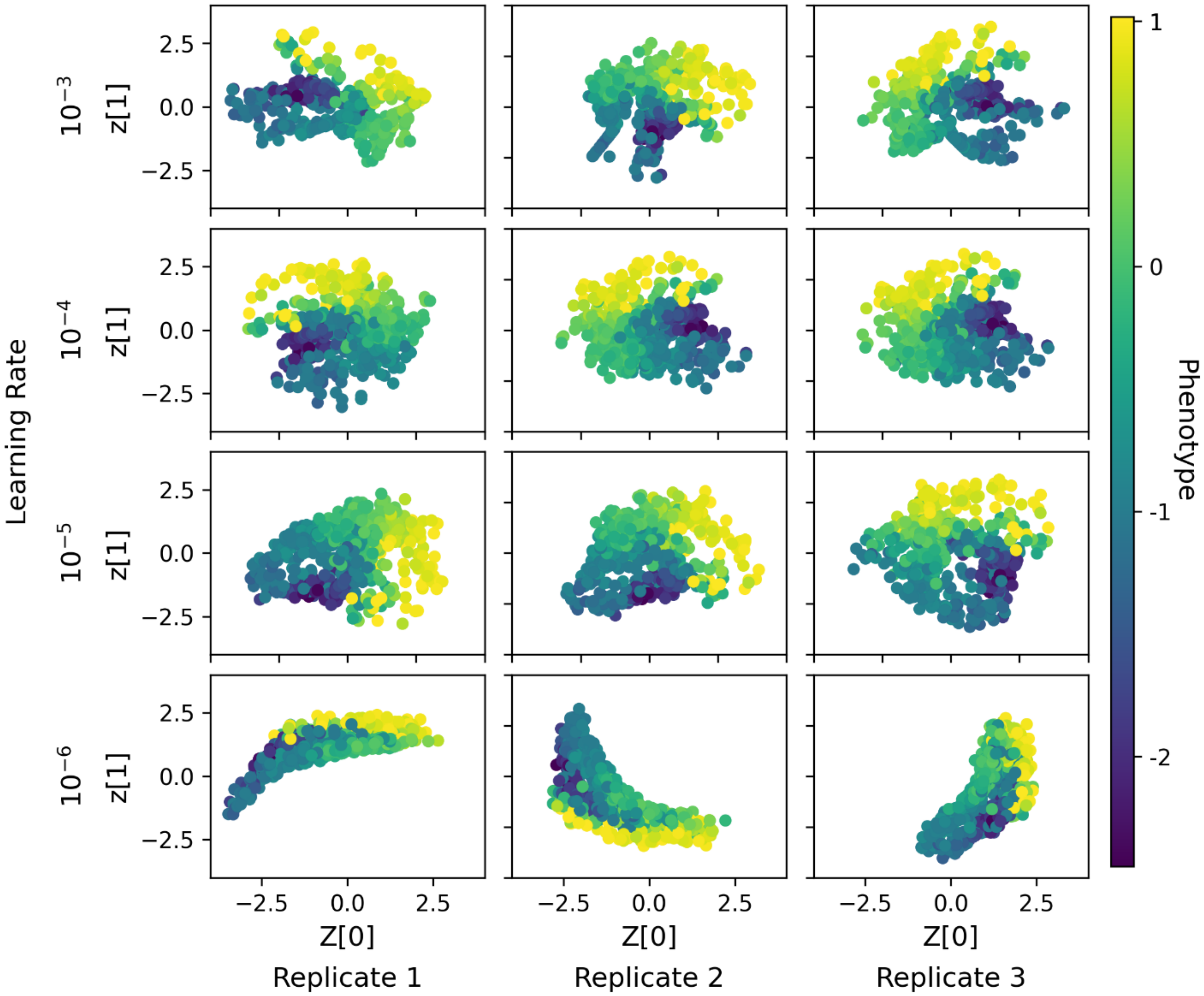


**Figure S16.** Three replicates of VAE models using the fitness data as features at different learning rates. Models with $10^{-6}$ learning rate were not converged. Models with $10^{-3}$ learning rate tend to be stuck in a local minimum loss. The models with $10^{-4}$ and $10^{-5}$ learning rates provided similar latent spaces without any convergence problem, therefore a learning rate of $10^{-4}$ was used for fitness-based models.

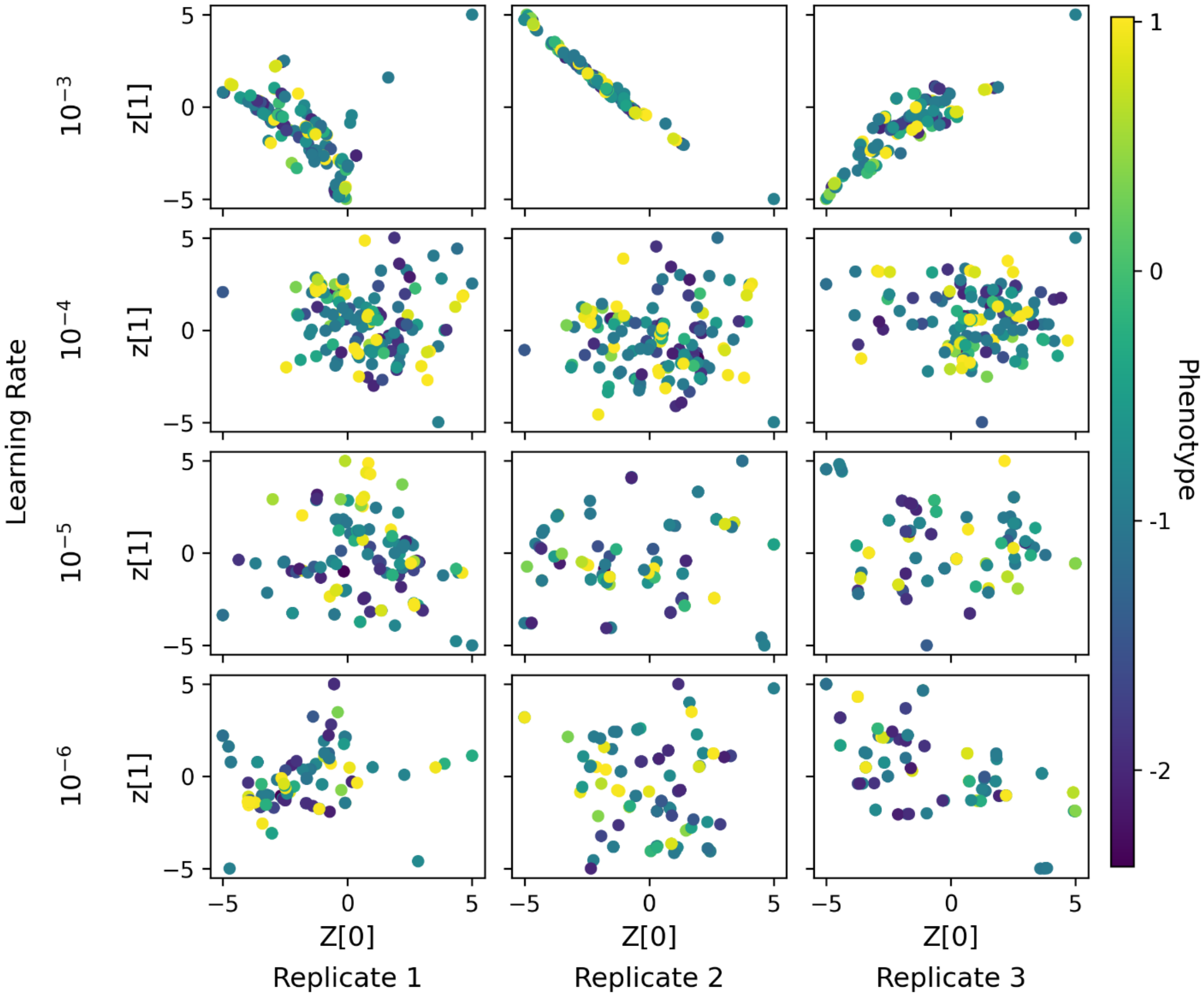


**Figure S17.** Three replicates of VAE models using the MD data as features at different learning rates. The same trend with the Fig S16 is observed. Learning rate of $10^{-4}$ provided the most visual separation of phenotypes, therefore it was used for the MD data models.

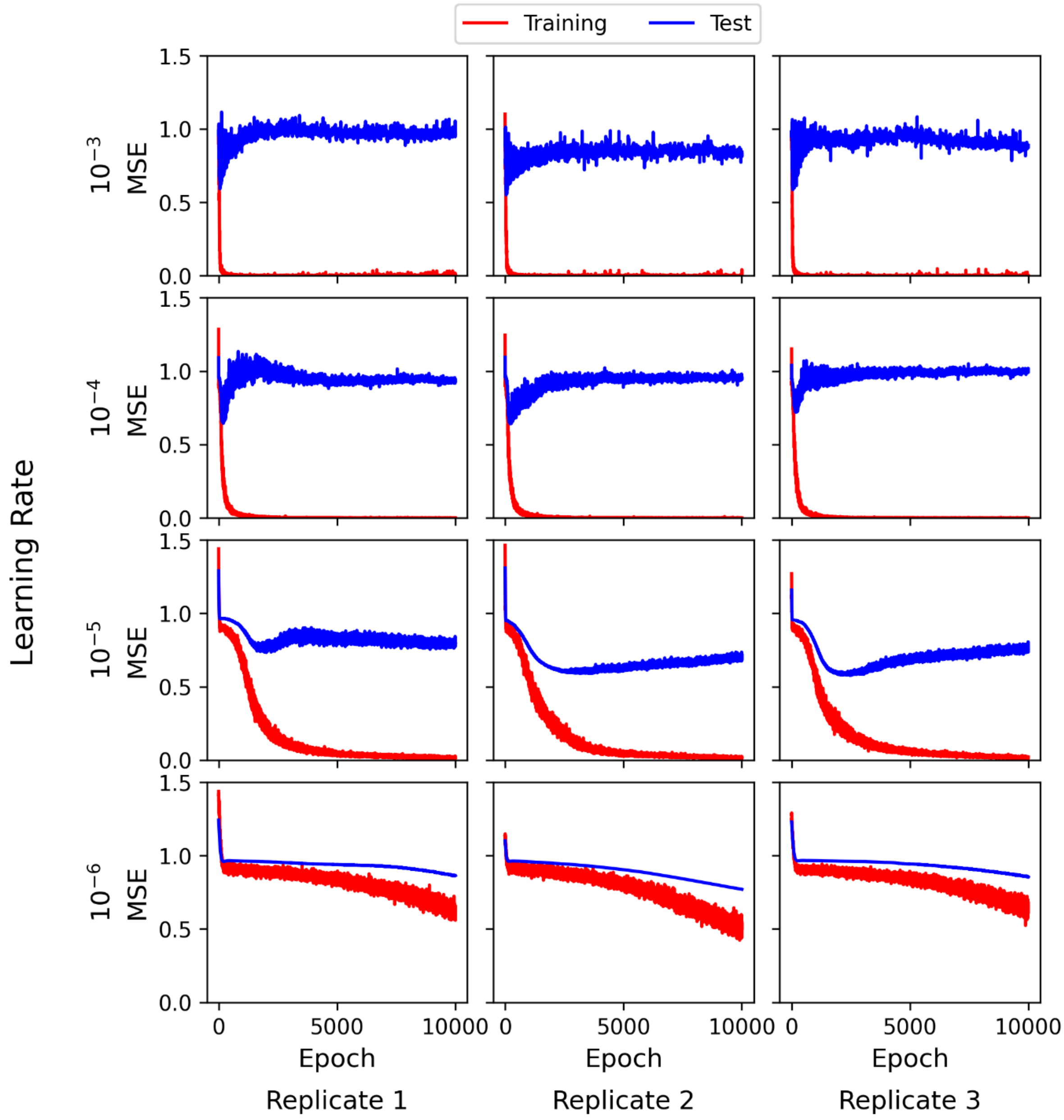


**Figure S18.** Three replicates of prediction models using the sequence data as features at different learning rates. Learning rate of $10^{-5}$ provided the minimum losses for the test sets, therefore it was used for the sequence models.

**Table S1.** The summary of deep learning models

| Model | Optimizer | Learning Rate | Epochs | Batch Number | Loss Function |
|---|---|---|---|---|---|
| Fitness – Prediction | Adam | $10^{-4}$ | 2000 | 4 | MSE |
| Fitness – VAE | Adam | $10^{-4}$ | 5000 | 4 | MSE+KL |
| Sequence – Prediction | Adam | $10^{-5}$ | 20000 | 100 | MSE+KL |
| MD – Prediction | Adam | $10^{-5}$ | 20000 | 100 | MSE+KL |
| MD – Prediction with Attention | Adam | $10^{-5}$ | 20000 | 4 | MSE |
| MD – VAE | Adam | $10^{-4}$ | 5000 | 4 | MSE+KL |

**Table S2.** Statistical analysis of the clusters obtained from VAE model using MD data. T-test was performed by assuming the two populations have different variance. Clusters I, II and III corresponds to the clusters shown in Figure 6A

| Metric | I | II | III | I-II | I-III | II-III |
|---|---|---|---|---|---|---|
| Mean | -0.49 | -0.75 | -0.83 | | | |
| Standard error | 0.11 | 0.26 | 0.14 | | | |
| T-statistics | | | | 0.90 | 1.96 | 0.29 |
| P-value | | | | 0.378 | 0.053 | 0.771 |
